# Semilichen an unjustly neglected symbiotic system between green biofilms and true lichens

**DOI:** 10.1101/2024.12.10.626781

**Authors:** Jan Vondrák, Stanislav Svoboda, Pavel Říha, Tomáš Hauser, Veronika Kantnerová, Pavel Škaloud, Jiří Kubásek

## Abstract

- Symbiotic systems of photosynthetic microorganisms and fungi are widespread in terrestrial biomes and lichens are probably the most advanced and complex. Conversely, the least complex systems are “green biofilms” with a completely unexplored mycobiome. We describe here a new system intermediate between green biofilms and lichens – semilichens.
- Light and fluorescence microscopy, eDNA sequencing, molecular phylogeny, Chlorophyll *a* fluorescence and ^13^C labelling/metabolomics were used to study algal and fungal identity, morphology and physiology of the symbiosis.
- Tight contact between algae and a single predominant fungus (mycobiont) is revealed in semilichens. The algae are from the symbiotic lineages of Trebouxiophyceae and Ulvophyceae, the fungi belong to Arthoniomycetes, Dothideomycetes, Eurotiomycetes, Lecanoromycetes and Lichinomycetes. Algae are alive and perform substantial photosynthetic activity. ^13^C labelled photosynthates are partially converted into specific fungal polyols (arabitol, mannitol) demonstrating the C-flow from algae to fungi.
- The new symbiotic system was defined and compared with other terrestrial algal-fungal symbioses. It is characterized by minimalist environmental requirements and extremely low production of biomass. As a result, it also inhabits environments unfavourable for lichens. Our research supports the hypothesis that the long-term existence of algae and fungi in terrestrial conditions affected by frequent and repeated drying is likely dependent on their mutual coexistence.

## Introduction

Symbiotic associations of fungi and algae are more common in terrestrial biomes than generally assumed, and curiously, some of the most common are not included in the current review on algal-fungal symbioses (Bonito, 2024). The largest omitted group are the so-called “green biofilms” (Freystein & Reisser, 2010). These are communities of aerophytic algae (sensu Ettl & Gärtner, 1995) which are widespread across most terrestrial biomes. Although these communities are generally dominated by algal biomass, fungal hyphae closely surrounding algal cells are almost always present (Fig. 4 in Freystein & Reisser, 2010; **Fig. 1, S1**, here). Green biofilms without fungi occur, according to our observations, only in habitats not subject to heavy periodic drying, e.g. at the base of tree trunks near the soil.

**Fig. 1.**
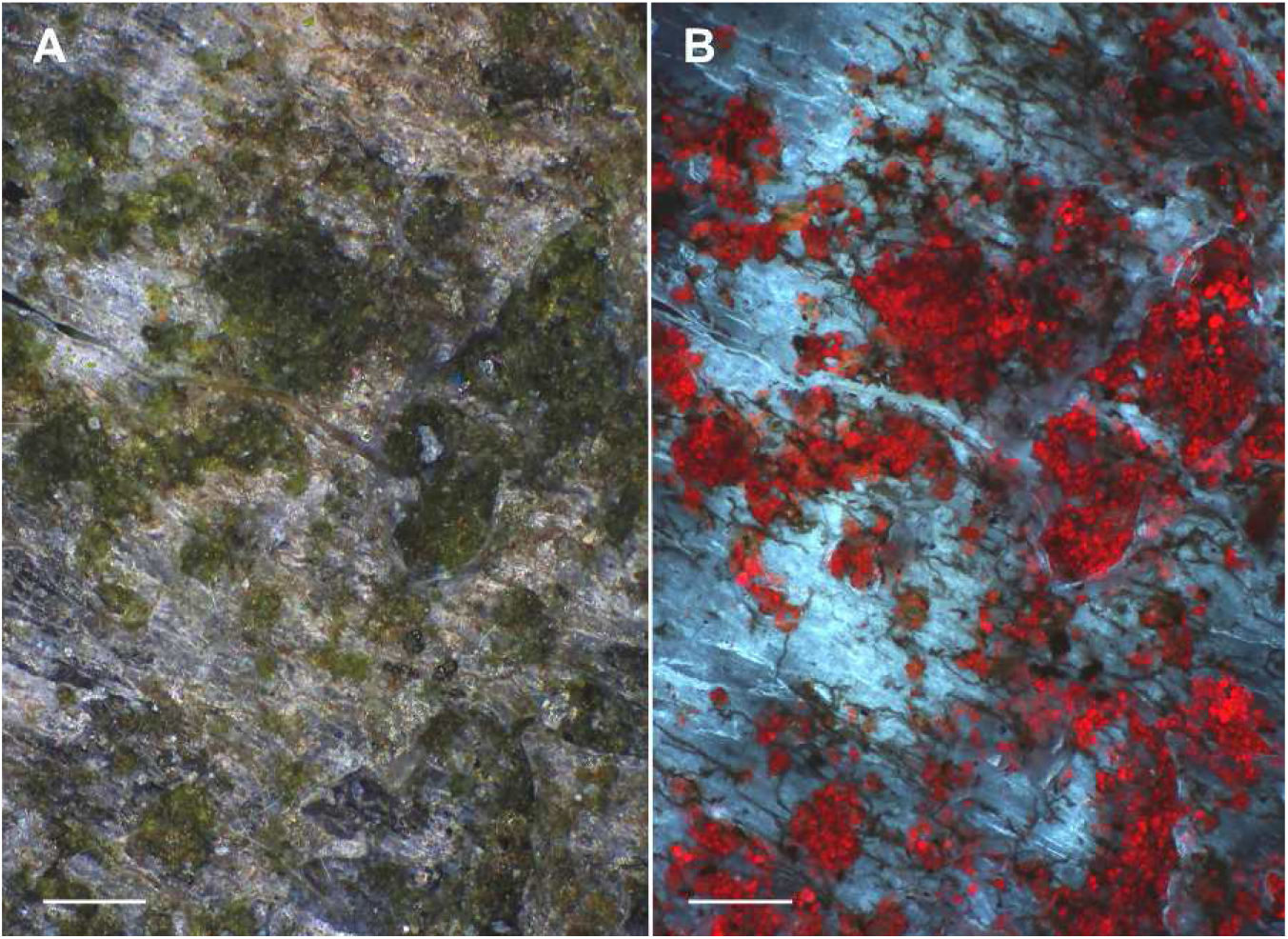
Corticolous green biofilm on bark of *Tilia* in urbanised landscape. A, observed in visible light where algal coating is conspicuous but the mycobiom is invisible. B, observed with fluorescence in blue light where the associated mycobiome is well recognised. Scales, 0.5 mm.

Another large group of algal-fungal symbioses, generally overlooked by science, are the semilichens. Although these symbiotic systems were introduced to the scientific community more than a century ago (as Halbflechten; Zukal, 1891), it is unlikely that a single study has been devoted to semilichens since then. Semilichens represent a kind of intermediate stage between green biofilms and lichens. Their basis is a single dominant fungal species (mycobiont), which produces specific fruiting bodies and thus the individual species of semilichens can be recognized by classical taxonomy. Semilichens, or more precisely their mycobionts, have long been recognized and studied by mycologists and lichenologists, usually as non-lichenised or facultatively lichenised fungi occurring in habitats together with lichens. They differ from true lichens in the absence of a conspicuous and stratified thallus with a recognizable algal layer. Therefore, the prevailing current opinion is that they are saprophytic/endophytic, non-lichenised fungi. The close coexistence of mycobiont with algae is however evident in semilichens, but it takes place in spatially separated colonies of algae.

It is possible that the close co-occurrence of algae and fungi is a necessary condition for the existence of these organisms in semilichens and biofilms on the surface of plants (epiphytes) or on the surface of rocks and stones (epilithes), which are exposed to extreme fluctuations in life conditions, especially frequent desiccation. The anhydrobiotic model developed by Spribille et al. (2022) represents an evolutionary hypothesis for the origin of algal-fungal symbiosis (specifically lichens) under terrestrial conditions based on the assumption that fungi, in order to expand their ecological niche by appearing “on the surface”, were forced to enter into a long-term symbiotic relationship with algae, which provide the fungi with substances necessary to survive the constant alternation of moisture and complete desiccation. A large group of such substances are polyols, which are widespread (Honegger, 1993; Gustavs *et al*., 2011). In contrast to sugars, polyols are more stable molecules employed in long-term carbon storage and osmoprotection (Gustavs *et al*., 2010). Aerophytic algae produce several polyols: glycerol (C3), erythritol (C4), arabitol (C5), mannitol (C6) and rarely volemitol (C7) in Trentepohliales, Ulvophycae (Feige & Kremer, 1980), ribitol (C5) in most Trebouxiophycae (Lewis & Smith, 1967) and sorbitol (C6) in Prasiolales and the Botryococcus-clade of Trebouxiophycae (Gustavs *et al*., 2011). Fungi associated with algae in terrestrial habitats utilize algal sugars and polyols (typically ribitol) and convert them into arabitol and mannitol (Richardson *et al*., 1967). Arabitol, with fast turnover, may be important for respiration/growth and mannitol, with slow turnover, may be used as osmoprotectant to survive fungal anhydrobiosis (hybrid model in Spribille et al., 2022). According to our observations, the anhydrobiotic model (elaborated further at the end of the Discussion) is applicable to most aerophytic algal-fungal associations, including semilichens and green biofilms, and is relevant to algae as well as fungi. Algae, originally aquatic organisms, expanded their range to terrestrial habitats, where wetness and dryness alternate, apparently thanks to coexistence with fungi.

In this study, we provide a definition of semilichens, and we describe several aspects of their biology: (1) phylogenetic identities of the algae and fungi involved, (2) morphology, especially the close coexistence of the two symbionts, (3) ecological aspect, (4) viability and photosynthetic activity of associated algae and (5) carbon transfer from symbiotic algae to the mycobiont.

## Materials and methods

### Sampling and morphological observations

Green biofilms are almost ubiquitous, and the studied material was collected from bark and stones in the urban landscape of České Budějovice (Czech Republic) in 2023. Observations of fungi and algae were made by light microscopy and fluorescence microscopy. Semilichens have been studied for a long time by the first author and the studied material comes from different places in the Czech Republic from the years 2022–2024 and is deposited in the herbarium PRA. Observations of semilichens were performed by light microscopy in combination with fluorescence in blue light and UV excitation. Samples were fixed in lactoglycerol cotton blue prior to microscopy to enhance negative staining (fluorescence absorption) of otherwise often colourless hyphae. Furthermore, red chlorophyll autofluorescence in UV and blue light was used for visualization. Semilichens were observed mainly on the surface of the substrates, but in some cases the substrate (i.e. usually young smooth bark of trees) was subjected to horizontal sections in order to observe the occurrence of hyphae and algae also inside the substrate.

### DNA metabarcoding and bioinformatics

DNA of the semilichen samples was isolated using a cetyltrimethylammonium bromide (CTAB)-based protocol (Aras & Cansaran, 2006). ITS2 rDNA amplicons were produced by PCR with the barcoded primers ITS3 (5’-GCA TCG ATG AAG AAC GCA GC-3’) and ITS4 (5’-TCC TCC GCT TAT TGA TAT GC-3’). The PCRs were performed using the Q5 High-Fidelity DNA polymerase (BioLabs Inc.), they were run in 35 cycles and the conditions were: initial denaturation at 98 °C for 30 s, 98 °C denaturation for 10 s, 52° C amplification for 45 s and 72 °C elongation for 1 min, with a final 72 °C extension for 2 min. Each sample was run in two replicates, and both negative controls (distilled water as a template) and multiplexing controls (unused combinations of left and right barcodes) were used. The PCR products were purified with SPRI AMPure XP paramagnetic beads (Beckman Coulter), pooled equimolarly and sent for library preparation and sequencing to Fasteris (Plan-les-Ouates, Switzerland). Sequencing was performed on the Illumina MiSeq platform with paired end mode (2 × 300 bp). Quality control of the Illumina MiSeq paired-end reads was carried out using FastQC v. 0.11.8 (Andrews S, 2010). Raw reads were processed according to Bálint et al. (2014), including quality filtering, paired-end assembly, removing primer artifacts, extracting reads by barcodes, reorienting reads to 5′-3′, demultiplexing, dereplicating, OTU clustering (this step carried out using Swarm v. 2 (Mahé *et al*., 2014), with denoising set to d = 3) and chimera filtering. Only swarms that were found in at least 10 reads in both replicates, and that were absent from the negative controls, were considered. The data are available at: http://www.ncbi.nlm.nih.gov/bioproject/PRJNA1184125. The OTUs were identified by BLAST searches in SEED2 (Větrovský *et al*., 2018), and only Viridiplantae and Fungi sequences were further processed. The obtained algal sequences were aligned with the closest BLAST matches and a maximum likelihood (ML) tree was constructed in IQ-TREE v. 1.6.1 (Nguyen *et al*., 2015) using GTR+I+G substitution model and ultrafast bootstrapping with 2,000 replications. Monophyletic clades were assigned to individual species based on historically defined species boundaries for each genus. Morphological identifications of mycobionts were confirmed by the Blast search against NCBI.

### Chrorophyll a fluorescence imaging

FluorCam FC 800-C (Photon System Instrument, Drásov, Czech Republic) was used for 2D fluorescence imaging. Pulse amplitude modulation (PAM) fluorescence was used. Measuring light averaged intensity was below 1 µmol m^-2^ s^-1^, pulse duration 20 µs, actinic light intensity 150 µmol m^-2^ s^-1^ and saturating pulse intensity 1000 µmol m^-2^ s^-1^ . Chlorophyll fluorescence was used to estimate: (1) chlorophyll abundance and distribution within sample, (2) Maximum quantum yield (F_v_/F_m_), which is a measure of algal viability/activity and (3) kinetics of photochemical quenching, which is a measure of relative photosynthesis rate. Plant twigs bearing semilichens, however, contain substantial amount of chlorophyll in their tissues (particularly in pheloderm layer). Via gentle drying for several days and subsequent rewetting, plant tissues were killed (F_v_/F_m_ close to zero) and surface algae resurrected (F_v_/F_m_ of healthy phototrophs range about 0.6 to 0.8). Thus, algal colonies were easily visible on the plant surfaces and their physiology could be further examined. In next step, we removed symbiotic algae from a part of the plant surface using a scalpel and compared it with native sample.

### ^13^C-transfer from algal to fungal polyols

This study uses isotope labelling, which has been used successfully in alcobioses (Vondrák *et al*., 2023) and much earlier in classical papers on lichen physiology (Richardson & Smith, 1968). Briefly, the stable carbon ^13^C, which behaves physiologically the same as widespread ^12^C, is scarce in nature. Thus, it is a sensitive label of new assimilate production and its fate. We added ^13^CO_2_ (99 At%, Cambridge isotope laboratories) into our labelling device used previously (Kubásek *et al*., 2021; Vondrák *et al*., 2023) to reach about twice atmospheric CO_2_ concentration (≈ 850 µmol CO_2_ mol^-1^) and let algae/symbioses assimilate it under mild light (200 µmol of photons m^-2^ s^-1^) for one or two hours. After that, part of the biological system (about 100 mg of fresh weight representing 0.5 to 2 cm^2^ patches of more or less planar symbiotic systems) was removed from the substrate, using a scalpel, and quickly killed in 2 mL of boiling methanol and dichloromethane (1:1) mixture. The rest of the system was left in normal conditions to translocate/convert assimilates from algae to fungi or to different metabolites (so called “chase period”), and harvested thereafter (see results).

Then, the supernatant was filtered, evaporated under nitrogen stream and OH groups of sugars and polyols silylated using 25 µL of BSTFA in 100 µL of pyridine and heated at 85°C for one hour. Finally, n-hexane was added to reach a total sample volume of 1 mL. Compound specific isotope analysis was performed using gas chromatography (Trace 1310, Thermo Scientific, Bremen, Germany) and Isotope ratio mass spectrometry (Delta V Advantage, Thermo Scientific, Bremen, Germany). Using chromatographic column ZEBRON ZB-1 (30 m x 0.25 mm ID and 0.25 µm film thickness) we were able to separate all principal polyols (erythrytol, ribitol, arabitol, manitol, sorbitol) and main sugars (glucose and saccharose).

We calculated the ^13^C excess as a difference between the ^13^C content of the sample/compound and its natural abundance (1.07 ± 0.005 At%). The relative amount of new carbon in a particular compound may thus be estimated as the ^13^C excess in that particular compound multiplied by the abundance of this compound in the sample and standardised to 100 % for sum of all compounds.

However, this relative partitioning of new carbon tells us nothing about the absolute rate of CO_2_ assimilation (a measure of whole system activity). Therefore, we calculated the average ^13^C excess in the pool of all compounds. In short, this is the amount of ^13^C (above its natural abundance) in the “average metabolite”. Since nearly pure ^13^CO_2_ is used, the 1% excess of ^13^C implies that 1 % of all mobile carbon has been replaced due to our labelling. The drop in this value one and four days after labelling represents both respiratory loss and incorporation of new carbon into structural polymers no longer accessible to gas chromatography. Owing to the abovementioned low variability of natural ^13^C content and high precision of isotope ratio mass spectrometry, the appearance of new carbon in particular compounds is well confirmed. On the other hand, physiological/ecological significance should be considered critically (see Discussion).

## Results

### Definition of a semilichen

A symbiotic system formed by a single dominant fungal species (mycobiont), living in facultative or obligate symbiosis with one or more species of algae. It does not form a three-dimensional and stratified thallus with a recognizable algal layer, as real lichens do, but the close coexistence of mycobiont and algae takes place inconspicuously in spatially separated spots, i.e. algal colonies. This is reflected in the minimalism in biomass production; lichens and green biofilms usually produce noticeably more biomass per area. The algae do not form a conspicuous coating and their biomass does not predominate over that of fungi, which makes semilichens markedly different from green biofilms. Hyphae of semilichen mycobionts are mostly exposed to the surface or are slightly immersed, but do not deeply penetrate the substrate. It is a long-term coexistence in which the algae thrive (so it is not an obvious parasitism of the fungus on the algae). Metabolic exchange between the algae and fungus occurs. The ecology partially overlaps with lichens and green biofilms, but semilichens can inhabit extreme microsites where lichens and green biofilms are at a strong disadvantage (see ecological section below). As with lichens, the name used informally to refer to a species of semilichen is the name of the mycobiont, although, strictly speaking, that name refers only to the mycobiont.

### Phylogenetic perspective

Semilichen mycobionts are present within the Ascomycota in Arthoniomycetes, Dothideomycetes, Eurotiomycetes, Lecanoromycetes (only in Ostropomycetidae) and Lichinomycetes (sensu Díaz-Escandón et al., 2022). According to our long-term observations in Europe, semilichen mycobionts are found in the following taxa:

### Arthoniomycetes

*Arthonia* and *Naevia*. **Dothideomycetes:** *Alloarthopyrenia, Arthopyrenia, Cyrtidula, Julella, Lichenothelia, Melaspilea, Melaspileella, Mycoporum, Naetrocymbe* and *Tomasellia*. **Eurotiomycetes:** *Atrodiscus, Blastodesmia, Chaenothecopsis, Leptorhaphis, Mycocalicium, Phaeocalicium* and *Stenocybe*. **Lecanoromycetes, Ostropomycetidae:** *Absconditella, Cryptodiscus, Eopyrenula, Epigloea, Exarmidium inclusum, Karstenia, Microcalicium, Sphaeronema truncatum* and *Xerotrema*. **Lichinomycetes (s.lat.):** *Steinia* and *Thelocarpon*.

Semilichen algae (photobionts) were identified by DNA barcoding from 14 symbiotic systems studied in detail (**Fig. 2**). These systems were generally dominated by algae from the Trebouxiophycae and often by species known from lichen symbioses (e.g. *Apatococcus lobatus, Elliptochloris antarctica* or *Trebouxia gelatinosa*). More rarely, the systems were dominated by algae of the genus *Trentepohlia* (Ulvophycae). A total of 25 algal OTUs were found (only species abundant in at least one sample are counted) from genera *Apatococcus* (4 OTUs), *Coccomyxa* (2), *Elliptochloris* (1), *Symbiochloris* (5), *Trebouxia* (9), *Trentepohlia* (3), and *Tritostichococcus* (1).

**Fig. 2.**
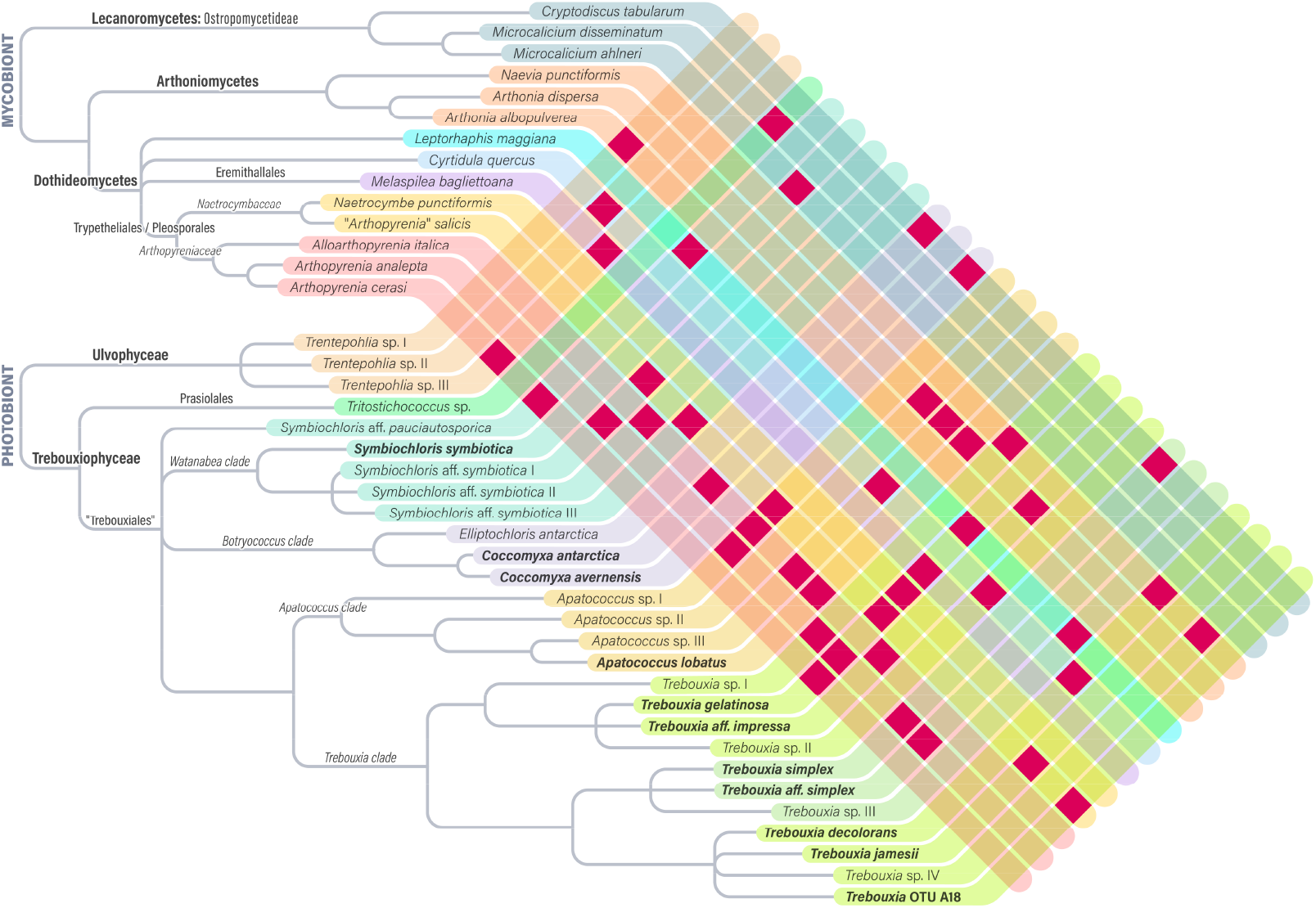
Fungal and algal systematic units involved in the fourteen examined semilichens. Algal species/OTU’s previously known from lichen symbioses are in bold. Specific relationships between mycobiont species and algal species/OTUs are illustrated by the dark coloured squares.

### Morphological observations of algal-fungal co-existence

Detailed morphological observations were made on the semilichens *Arthopyrenia analepta* (**Fig. 3**), *A. cerasii* (**Fig. 4**), *A. salicis* (**Fig. S2**), *Cryptodiscus tabularum* (**Fig. S3A**,**B**), *Cyrtidula quercus* (**Fig. S4**), *Karstenia idaei* (**Fig. S3C**,**D**), *Naetrocymbe punctiformis* (**Fig. 5**) and *Naevia punctiformis* (**Fig. S5**). In most cases, a significant part of the fungal hyphae is exposed to the external environment on the substrate surface. These hyphae are associated with algae that occur on the surface and usually are clustered in scattered colonies (**Fig. 3, 5**).

**Fig. 3.**
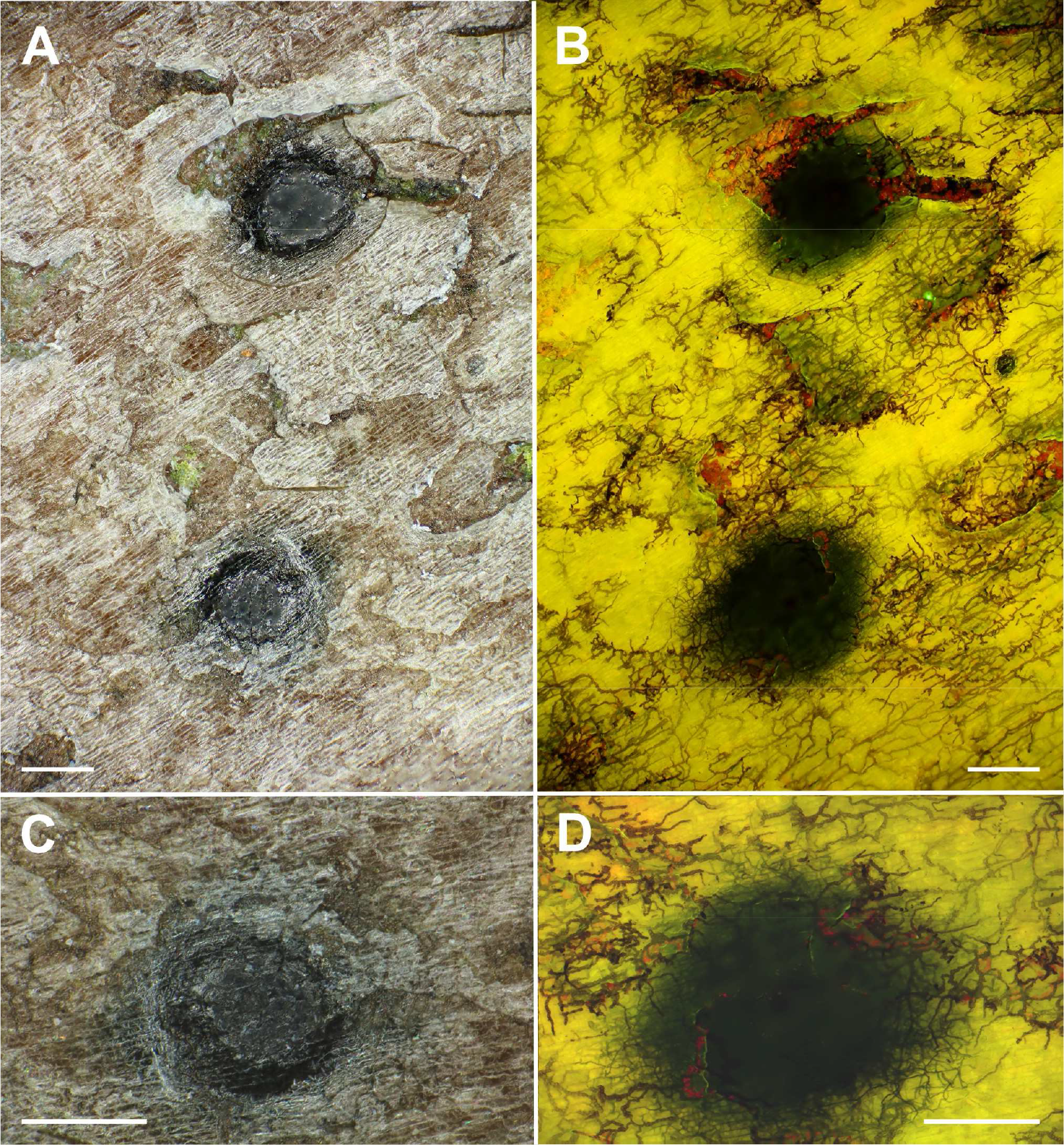
Semilichen *Arthopyrenia analepta* (Dothydeomycetes, Trypethelliales) with perithecioid fruiting bodies. A, C, observed in visible light, hyphae of mycobiont invisible. B, D, observed with fluorescence in blue light where the algal-fungal association is readily visible. Scales, 0.2 mm.

**Fig. 4.**
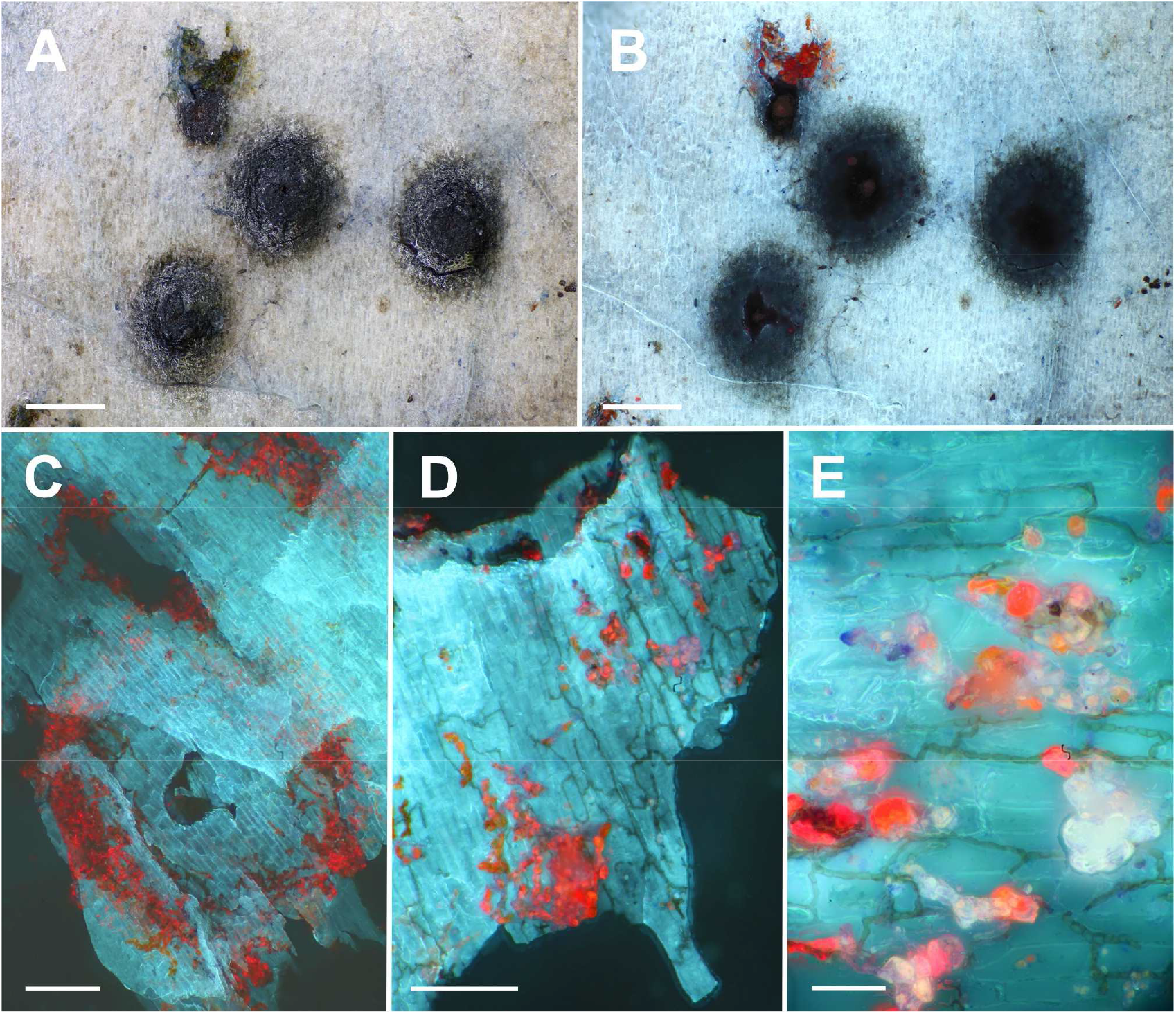
Semilichen *Arthopyrenia cerasi* (Dothydeomycetes, Trypethelliales). Algal cells occur under a layer of smooth scaly bark and are invisible by observation of the surface. A, B, fruiting bodies (perithecia) observed on the surface of bark of *Corylus avellana*. A, observed in visible light; B, observed with fluorescence in UV. C–E, algal-fungal association observed on a bark layers several tens of micrometres below the surface. Scales, A, B, 0.2 mm; C, D, 50 µm; E, 10 µm.

**Fig. 5.**
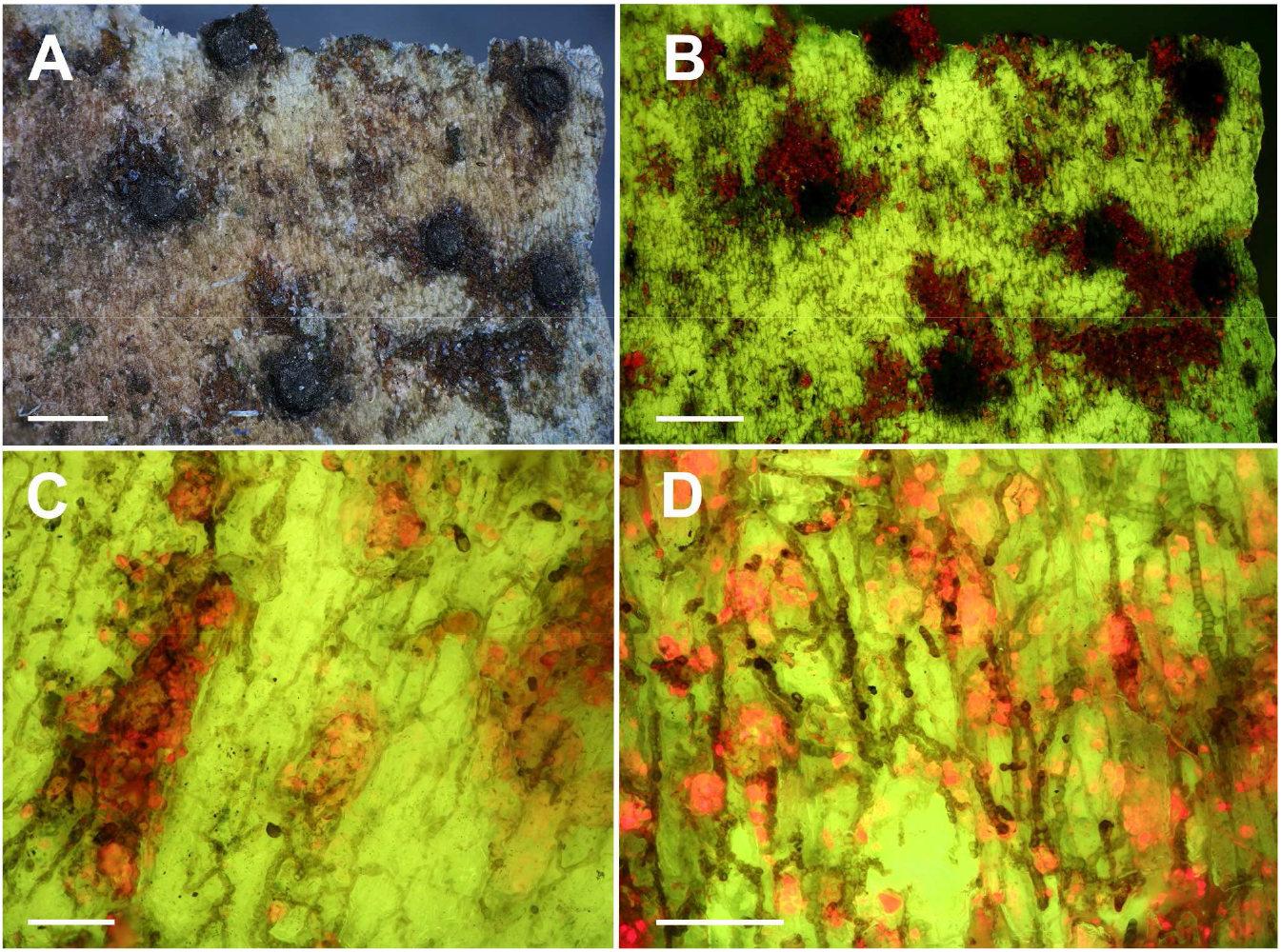
Semilichen *Naetrocymbe punctiformis* (Dothydeomycetes, Capnodiales). A, B, fruiting bodies (perithecia) on the surface of bark. A, observed in visible light, hyphae of mycobiont invisible; B, observed with fluorescence in blue light, fungal hyphae surrounding algal colonies readily visible. C, D, details of algal-fungal association with fluorescence in blue light. Scales, A, B, 0.2 mm; C, D, 50 µm.

In some cases, the hyphae network is spread beneath the surface, under a layer of smooth scaly bark several tens of micrometres thick, and is thus invisible by simple surface observation (**Fig. 4A**,**B**). However, if we observe in an appropriate layer below the surface we find similar structures as in semilichens with hyphae on the surface (**Fig. 4C-E**). It is similar to the hypophloedal thallus of some lichens, but here the algae do not form a conspicuous continuous layer.

Fungal hyphae are either colourless or melanized to varying degrees. In both cases, they are difficult to observe with classical light optics (**Fig. 3-5**), which is probably why the symbiosis of semilichens has escaped attention for so long. However, mycobiont hyphae are readily observable by fluorescence microscopy, especially after cotton-blue staining, as they absorb blue light and UV (false dark coloration), contrasting with the distinctly paler surrounding substrate.

### Ecology – semilichens occupy open niches

Most known semilichens are epiphytes, only a small proportion occur on rocks or stones (e.g. genus *Lichenothelia*) and some are part of soil microbial crusts (*Steinia, Thelocarpon*). In this study we are mainly concerned with epiphytic semilichens which are widespread in Europe, both in natural habitats and in habitats heavily disturbed by humans. Their ecology is to some extent similar to lichens or corticolous green biofilms, but their niches often differ, and semi-lichens fill the space not occupied by other symbiotic systems.

Some semilichens find a specific niche on young substrates (sprouts, twigs), where more complex lichens are less competitive because of their slower development. The two most common semilichens in Central Europe are *Naetrocymbe punctiformis* and *Naevia punctiformis*, which primarily overgrow thin twigs with smooth bark, but in heavily acidified areas with impoverished epiphytic lichen communities (i.e. in “lichen deserts”; (Gilbert, 1971), they often overgrow entire tree trunks, replacing lichen communities.

Other semilichens are able to grow in conditions that are too shady to allow lichens to grow, or only to a limited extent. For example, *Cyrtidula quercus* and *Leptorhaphis maggiana* have been observed on rods of *Corylus avellana* in very dark forest conditions. Semilichens are also often able to grow in micro-sites that are too dry and sunny, where lichens cannot grow. An extreme case is *Microcalicium loraasii*, which grows on sunny and parched conifer bark. Life in extreme conditions is probably made possible for semilichens by the minimalist lifestyle and the low requirements of the low-biomass partners.

### Algae thrive in semilichens

The maximum quantum yield of PSII (F_v_/F_m_), powerful indicator of photosynthetic apparatus activity/viability, shows that surface dwelling symbiotic algae are viable, contrasting with green plant tissues (pheloderm), particularly after drying and rewetting (**Fig. 6**). Despite the low abundance of algal chlorophyll in comparison to green plant tissues, we can see F_v_/F_m_ about 0.6 in rewetted algal colonies in contrast to near zero in the dead plant tissues (**Fig. 6**, right column). These values for semilichens (**Fig. 6F, I, L**) are comparable to algae in green biofilms (**Fig. 6C**) and in true lichens (**Fig. 6O**), although chlorophyll abundance is clearly highest in the true lichen *Graphis scripta* (the largest area with high F_v_/F_m_). F_v_/F_m_, however, indicates the potential for photosynthesis, but not the actual rate of photosynthesis. Thus, we measured also chlorophyll fluorescence kinetics (Kautsky induction) in the semilichen *Cyrtidula quercus* (**Fig.7**), a green biofilm, the semilichen *Arthonia salicis* and the true lichen *Graphis scripta* (**Fig. S6**). The upper plot in each graph demonstrates the visual situation and selected pixels for analysis (separately: symbiosis, green plant tissue and calibration plate), middle plot is absolute value kinetics and bottom plot shows normalised fluorescence kinetics to F_o_ = 1. We see a comparable decline of fluorescence (=increase of photosynthesis) between 20 and 90 s where actinic light (= light driving the photosynthesis) was applied in all algal-fungal systems but not in killed plant tissue nor calibration plate. Algae in semilichens, green biofilms and true lichens express comparable photosynthesis activity, but are spatially scattered in semilichens (while omnipresent in true lichens and some green biofilms).

**Fig. 6.**
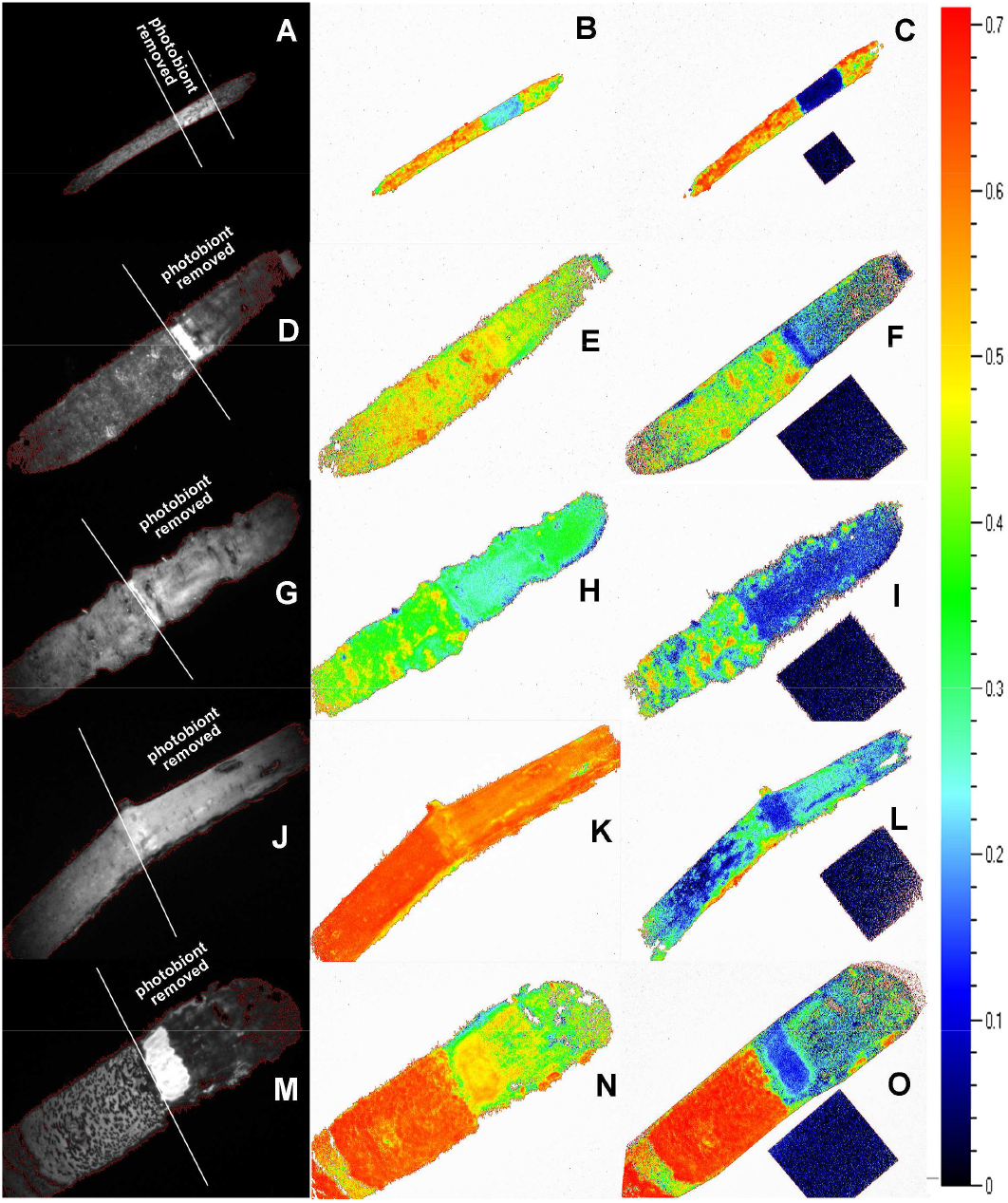
Maximum quantum yield of photosystem II (F_v_/F_m_) for a green biofilm (A-C), semilichens (D-L) and a true lichen (M-O). Left column shows fluorescence intensity, which is related to chlorophyll content. Photobionts were removed from part of samples using scalpel (areas delimited by white lines). The most intensive fluorescence is visible in patches where rhytidoma, masking green plant tissues, was also removed (especially D,M). Middle column are native samples, whereas right column are samples dried for four days in laboratory conditions to deactivate/kill green plant tissues and rewetted one hour before measurement again. Semilichens are: *Arthonia salicis* (D-F), *Cyrtidula quercus* (G-I) and *Naetrocymbe punctiformis* (J-L). True lichen is: *Graphis scripta* (M-O). Calibration plate (zero F_v_/F_m_) has dimension of 20 x 20 mm.

**Fig. 7.**
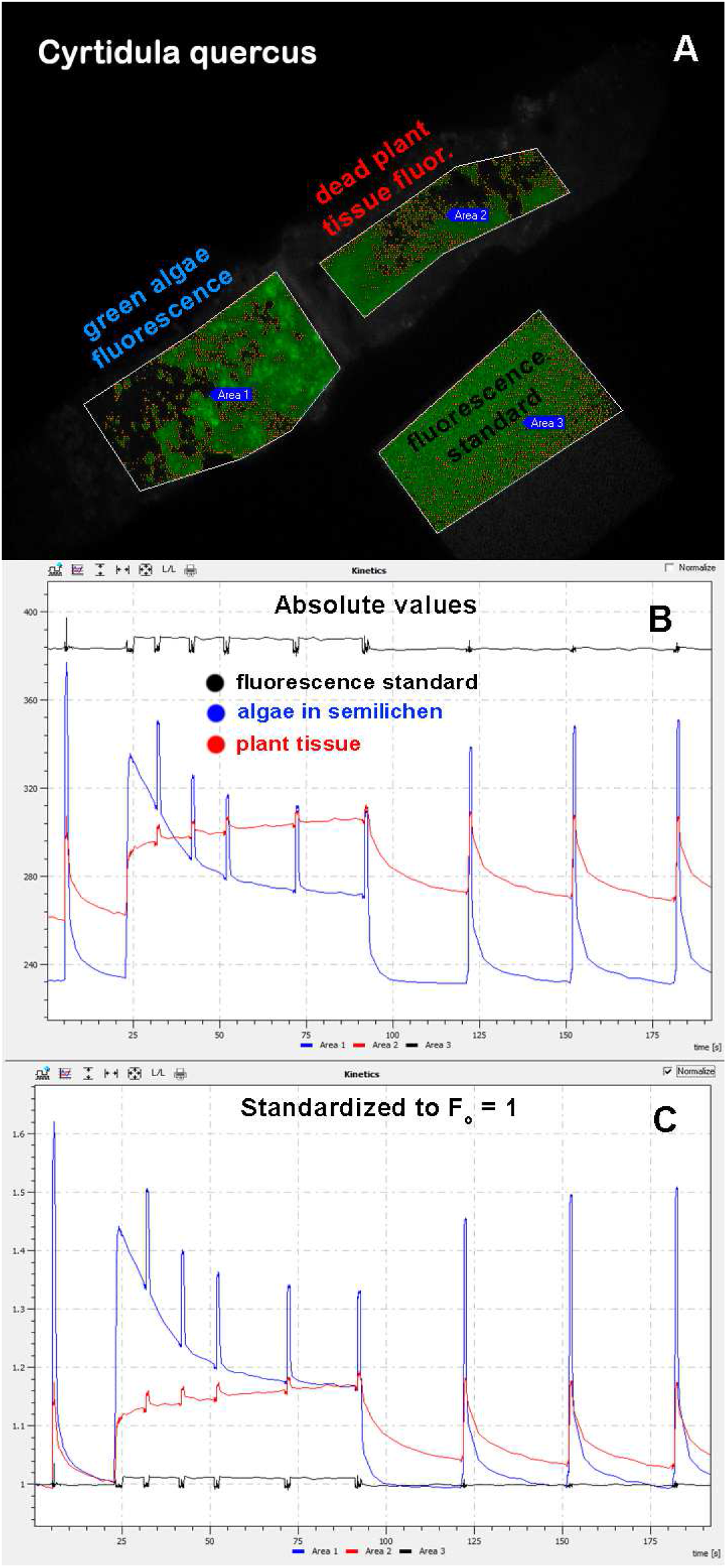
Chlorophyll *a* fluorescence kinetics of the semilichen *Cyrtidula quercus*. Visual situation (A), absolute values fluorescence (B) and fluorescence standardised to F_o_ = 1 (C). Comparison of symbiosis (blue line, area1), plant tissue where photobionts were removed (red line, area2) and calibration plate where no photochemistry occurs (black line, area3) is made. Dark adapted sample was subjected to low intensity measuring light (< 1 µmol m^-2^ s^-1^) for first 4 s, then to saturating flash (≈ 1000 µmol m^-2^ s^- 1^) for one second allowing us to calculate maximum quantum yield of PS II (F_v_/F_m_). After short dark relaxation (6 to 19s), in time 20 to 90 s, actinic light (150 µmol m^-2^ s^-1^) with five superimposed saturating flashes were applied to find photochemical (photosynthesis) and non-photochemical (photoprotection) quenching. Last part (91-190s) is dark relaxation with three saturating flashes to obtain relaxation rate of photoprotective mechanisms.

### ^13^C-transfer from algal to fungal polyols

Fractions of new carbon (^13^C) in different sugars and polyols at three times after labelling are shown in **Fig. 8** (green biofilm, semilichens) and **Fig. 9A-D** (true lichens). Owing to the specificity of particular polyols to algae or to fungi, putative carbon transfer between partners may be estimated. A slight but conclusive transfer of carbon from algal ribitol and sucrose to fungal mannitol and arabitol was observed in a green biofilm (**Fig. 8A**) and in examined semilichens (**Fig. 8B-F**). A similar pattern is visible for four true lichens (**Fig 9A-D**). Typically, the dominant fraction of new carbon is in ribitol and sucrose immediately after labelling (2h), and progressively moving into arabitol and mannitol during four-days chase period (assimilation in normal air). Some systems also produce substantial amount of four-carbon erythritol, which probably does not distinguish between algal and fungal pools (see discussion). The category “REST” represents all other GC-amenable metabolites, which were not identified.

**Fig. 8.**
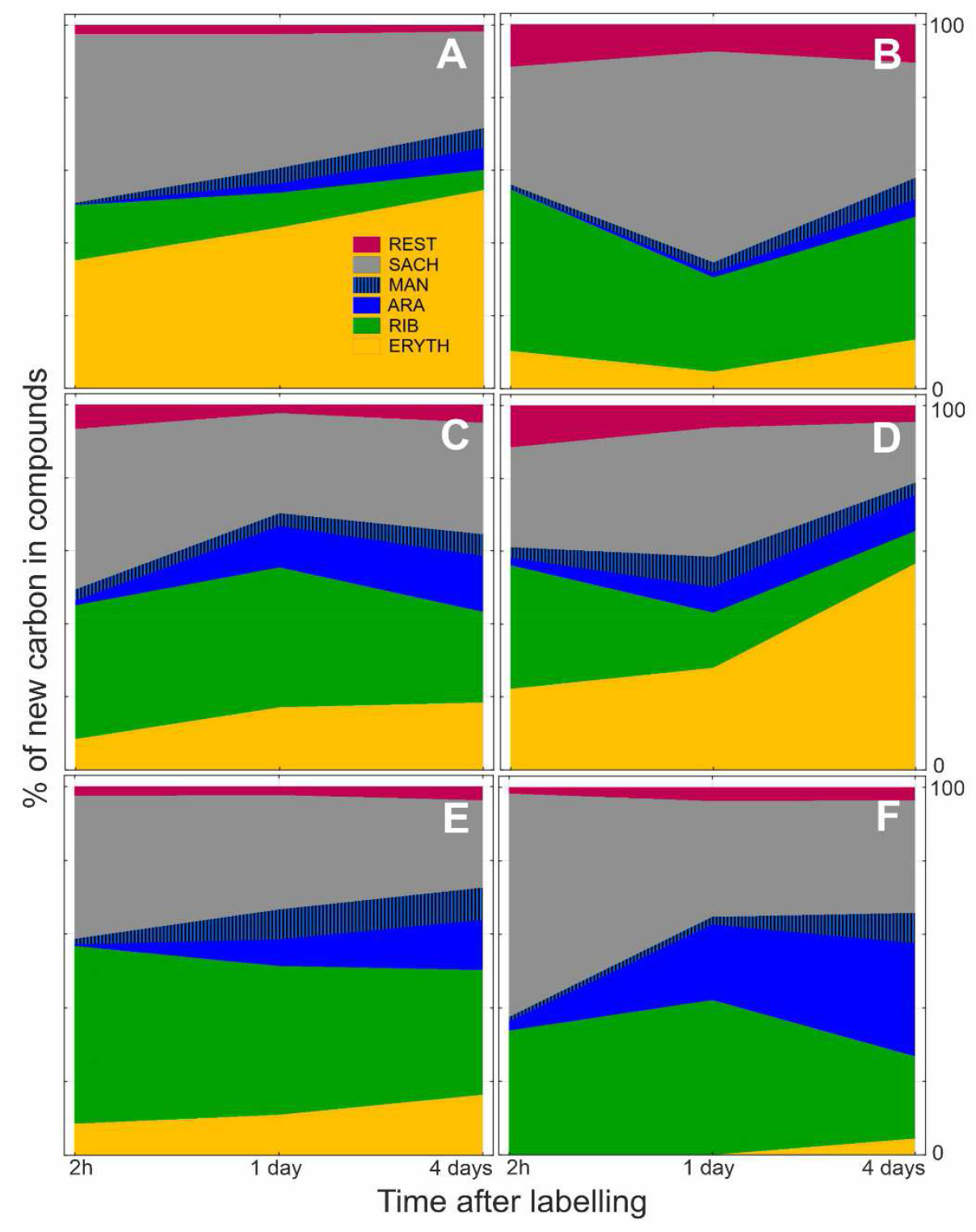
Green biofilm and semilichens – Percentage of new carbon in particular sugars and polyols 2 h, one day and four days after ^13^CO_2_ labelling. Gradual incorporation of new carbon into fungal polyols (arabitol and mannitol) is visible. A, corticolous green biofilm (see Fig. 1); B, C, two different samples of *Leptorhaphis maggiana* (Eurotiomycetes, Phaeomoniellales); D, *Naetrocymbe punctiformis* (Dothydeomycetes, Capnodiales); E, *Naevia punctiformis* (Arthoniomycetes, Arthoniales); F, *Stenocybe pullatula* (Ostropomycetidae, Mycocaliciales).

**Fig. 9.**
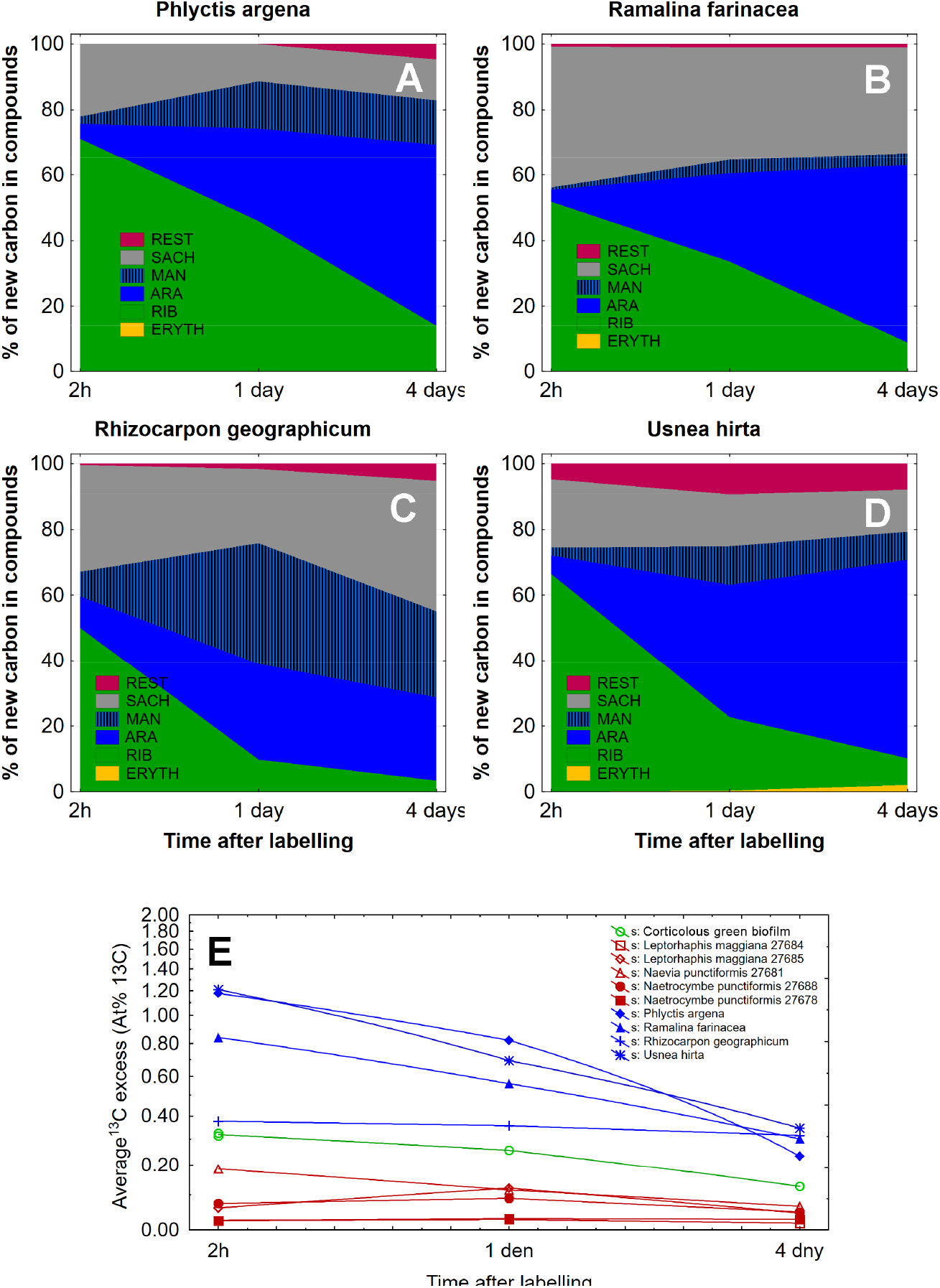
Lichens – Percentage of new carbon in particular sugars and polyols 2 h, one day and four days after ^13^CO_2_ labelling. Gradual incorporation of new carbon into fungal polyols (arabitol and mannitol) is visible. A, *Phlyctis argena* (Ostropomycetidae, Gyalectales); B, *Ramalina farinacea* (Lecanoromycetidae, Lecanorales); C, *Rhizocarpon geographicum* (Lecanoromycetidae, Rhizocarpales) and D, *Usnea hirta* (Lecanoromycetidae, Lecanorales). E, Average ^13^C enrichment in all metabolites pooled. It is measure of CO_2_ assimilation intensity (and reciprocally metabolite turnover) for whole symbiotic system.

Since we hypothesise that algal carbon (polyol) requirements are significantly lower in the small fungi in green biofilms and semilichens than in the biomass-rich true lichens, we calculated average ^13^C excess. Four exemplified true lichens reached higher ^13^C enrichment than five semilichens at all times after labelling, with green biofilm being intermediate (**Fig. 9E**).

All these and some additional systems are shown also in **Fig. S7** and **Fig. S8**. Apart from the percentage of new carbon in particular compounds (**Fig. S7**, right column), we present the fraction of these compounds in the total metabolite pool (**Fig. S7**, left column). By comparing the percentage of each compound and the percentage of new carbon in it, we can estimate the turnover rate of the respective compound (higher ^13^C percentage and lower percentage of substance in total metabolite pool - faster turnover). Average ^13^C excess of all systems studied is in **Fig. S8**.

In the case of symbiotic systems with *Trentepohlia*, it is difficult to trace the carbon flux from the alga to the mycobiont, as demonstrated in the semilichen *Arthopyrenia salicis* (**Fig. S7**). This is because the algae of Trentepohliales produce both mannitol and arabitol (Feige & Kremer, 1980), i.e. polyols that are exclusively fungal in symbiotic systems with other algae.

## Discussion

### Aerophytic algal-fungal associations

The only symbiotic system of aerophytic algae and fungi that has been generally accepted so far is lichens. We emphasize here that there are at least three other systems that can be distinguished from lichens (see below).

#### (1) Lichen (Fig. 10A)

A symbiotic system formed by a single dominant fungal species (mycobiont) and one or more species of algal or cyanobacterial photobiont. It is characterised morphologically by the existence of a three-dimensional “lichenised thallus” where the photobiont is internally organised, often in a layer. The lichen mycobiont depends on the supply of carbon, in the form of polyols and sugars, from its photosynthetic partner (Drew & Smith, 1967; Richardson *et al*., 1967). It is the best known and longest known coexistence of algae and fungi (Schwendener, 1869), and there are several recent reviews summarizing current knowledge (Lücking *et al*., 2021; Spribille *et al*., 2022; Pichler *et al*., 2023; Lücking & Spribille, 2024).

**Fig. 10.**
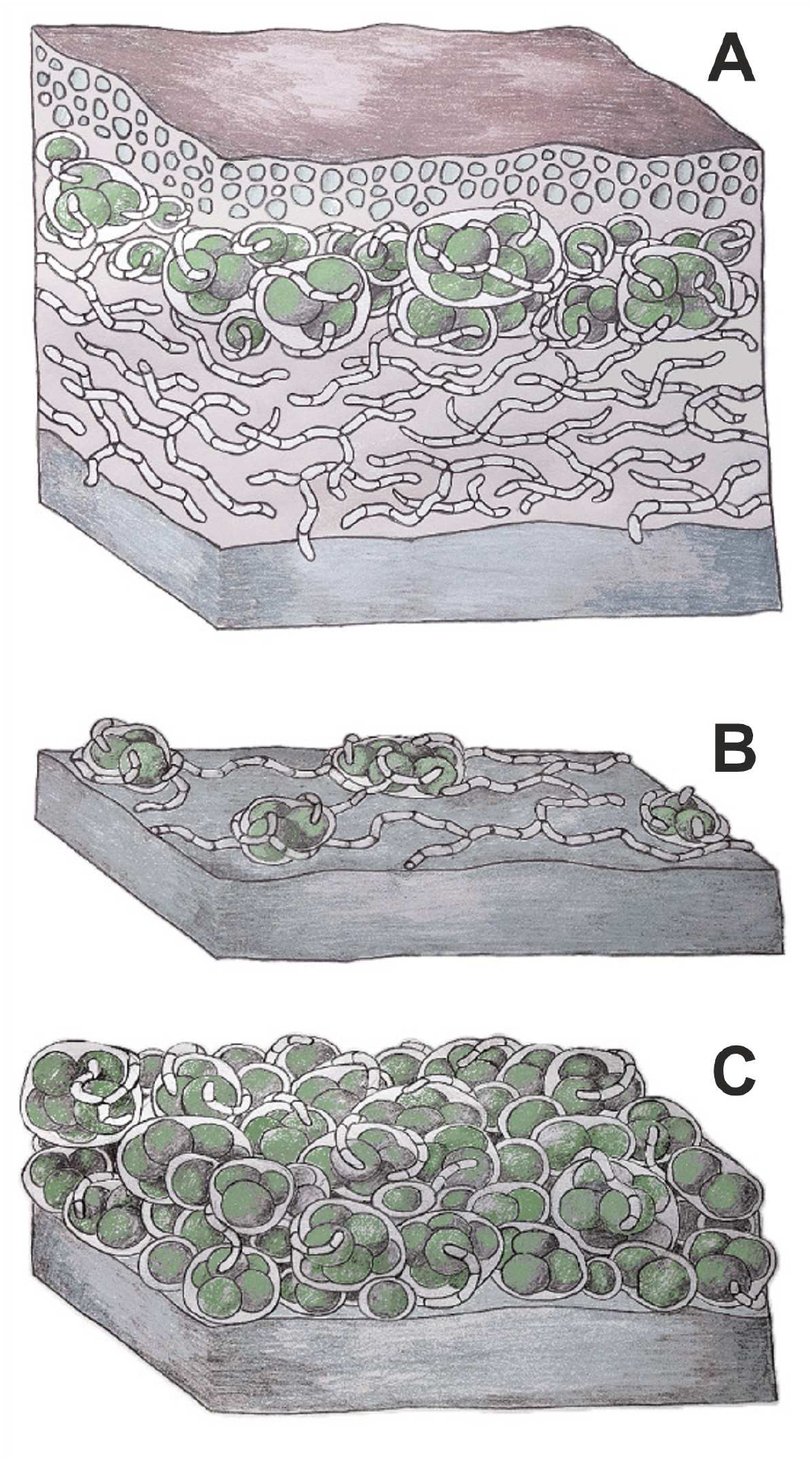
Schematic illustration of the most common aerophytic symbioses in temperate habitats; A, lichen; B, semilichen; C, green biofilm. The crustose lichen (A) is composed of a complex multi-layer thallus, consisting of upper cortex (isodiametric fungal cells), algal layer and medulla (loose hyphal tissue adjacent to the substrate). The semilichen (B) has a simple structure, consisting of a hyphal network growing over the substrate and associated algal colonies. The green biofilm (C) consists of aggregated algal cells and associated mycobiome.

#### (2) Semilichen (Fig. 10B)

The definition is given above in the results. The term was first used by Zukal (1891, as Halbflechten) for the minute fungi occurring on bryophyte leaves, which, according to his observations, always occurred in the company of algae. Unfortunately, the identity of the fungi that Zukal described in his article is now uncertain. Their names have fallen into oblivion and the type material, supposedly stored in a Viennese herbarium, has not been found. In the following more than a hundred years, no one was interested in Zukal’s concept, but taxonomic research on fungi, which we now consider to be mycobionts in semilichens, gradually proceeded. These fungi were thought to be non-lichenised or possibly associated with algae, but their relationship with algae has not been thoroughly investigated. The term “Halbflechten” has only recently been resurrected in its English form by (Vondrák *et al*., 2022), who, in their study on local lichen biodiversity, strictly distinguished semilichens, which are also briefly characterized as fungi associated with algae, but without the typical three-dimensional lichenised thallus. A more detailed characterization of the semilichens awaited our current study.

Some semilichens are hardly distinguishable from green biofilms. The algal coatings associated with e.g. the semilichen genera *Microcalicium* or *Stenocybe* have a very similar morphology to numerous corticolous green biofilms. Semilichens are not sharply delimited even from true lichens. For example, the symbiotic system of the fungus *Eremithallus costaricensis* (Lücking *et al*., 2008) represents an intermediate stage between a semilichen and a lichen.

A certain parallel to semilichens are the so-called borderline lichens occurring in the marine environment. They are symbioses of fungi with algae or cyanobacteria, which have a strongly reduced crustose thallus (Kohlmeyer *et al*., 2004; Pérez-Ortega *et al*., 2016).

#### (3) Green biofilm (Fig. 10C)

A community of aerophytic algae co-occurring with a wide range of microscopic fungi (mostly hyphomycetes). Algae form most of the biomass and are prominent at first sight, while fungi are inconspicuous due to their smaller biomass and often colourless or slightly melanised hyphae. The algae and fungi live in a close symbiotic relationship (**Fig. 1, S1**) and a metabolic exchange between them was demonstrated (**Fig. 7A**). Outwardly, the algal colonies co-occurring with the mycobiome are thriving and the fungi are not in the role of overt algal parasites, but rather in the role of long-term mutualistic symbionts. The relationship between microscopic algae and hyphomycete mycobiome is, along with the lichens, the most widespread algal-fungal relationship in terrestrial environments subject to periodic desiccation.

Microalgal communities in green biofilms on bark of trees are known to be diverse (Freystein & Reisser, 2010; Kulichová *et al*., 2014) consisting of green algae belonging mainly to Trebouxiophyceae and Chlorophyceae (Štifterová & Neustupa, 2015, 2017). While information on algal communities is rich, the surprising diversity of fungi and other groups, both prokaryotic and eukaryotic, is only recently beginning to be recognized (Freudenthal et al. 2024). Fungal hyphae closely attached to *Apatococcus lobatus* cells in green biofilms were reported by Geitler (1942) and Gärtner (1974). The furthest in mycobiome research was achieved by (Freystein & Reisser, 2010), who isolated axenic cultures of the alga *Elliptochloris* and accompanying hyphomycetes from a green biofilm and observed the development of these axenic cultures along with the development of the mixed community of fungus and algae. Their data showed that algae in the company of the fungus multiplied faster and lost chlorophyll more slowly, suggesting that the association with the fungus is beneficial to the algae. The taxonomic identity of fungi in green biofilms is still completely unknown.

In addition to algae, some biofilms also contain cyanobacteria. Recently, a specific symbiotic system of fungi and cyanobacteria, the phyllosymbium, has even been described, which, like semilichens, is intermediate between the lichen and the green biofilm (Chen *et al*., 2025).

#### (4) Alcobiosis

A hitherto little known but locally abundant symbiotic system in which algae form colonies or a continuous layer within and beneath the fruiting bodies of corticioid basidiomycetes (Vondrák *et al*., 2023). Morphologically, this coexistence bears a striking resemblance to lichens, but the dependence of the mycobiont on carbon from the algal partner has not been demonstrated and the benefits of this coexistence for both partners remain a mystery.

### Facultative lichens / facultative semilichens

The dependence of lichen mycobionts on carbon supply from their photosynthetic partners is probably not unequivocally true. At least some macrolichens, e.g. Peltigerineae and *Usnea*, have been shown to have laccases and the saprophytic enzyme activity (e.g. Laufer et al., 2009; Beckett et al., 2015) and they may therefore partly feed on organic matter from the substrate. Utilization of organic carbon from the substrate is even more likely in microlichens (not studied), and several species of fungi from the Ostropomycetidae are known that can associate with algae to form lichen, but can also exist without them (Wedin et al., 2004; Muggia et al., 2011). In our concept, therefore, such optionally lichenised fungi are not semilichens, but facultative lichens. In Ostropomycetidae, but also in other predominantly lichen lineages, some species congeneric with lichens are completely unlichenised (e.g. (Da Silva Cáceres *et al*., 2020).

In semilichens, the use of organic nutrients from the substrate is probably even more widespread, and some semilichen mycobionts can also occur almost without algae and can be called facultative semilichens (e.g. *Arthonia albopulverea, Naevia pinastri* or *Microcalicium ahlneri*). Some genera, in which semilichens predominate, also include representatives in which an association with algae has not been observed (e.g. *Naevia pinastri*; Thiyagaraja et al., 2020).

It is likely that the mycobionts of lichens and semilichens live part of their lives in the aposymbiotic phase. In true lichens, this phase seems to be mostly short and represents the spore germination stage, when the fungus seeks a suitable photosynthetic partner (Spribille *et al*., 2022; Pichler *et al*., 2023). In semilichens, however, it is possible that the aposymbiotic phase may be long-term and that the mycobiont behaves quite differently in this phase than in the symbiotic phase, e.g. as an endophyte, saprophyte or even parasite. This final suggestion is supported by the fact that relatively closely related fungi, such as those of the Dothideomycetes and Eurotiomycetes, exhibit a wide range of life strategies (Wijayawardene *et al*., 2020).

### The anhydrobiotic model is applicable to fungi in semilichens

The “nutritional model” of lichen symbiosis, prevailing until recently, assumed that the mycobiont uses the carbon obtained from its photosynthetic partner for respiration and growth (Smith, 1963). This model is supported by numerous data showing correlations between the photosynthetic activity of the phototroph and thallus growth (Palmqvist K *et al*., 2008). However, most of these data were obtained from “large” macrolichens, which are only the tip of the iceberg of the entire diversity of lichens and algal-fungal associations overall. Most lichens, however, have the form of small and slowly growing crusts, potentially more physiologically similar to semilichens.

(Spribille *et al*., 2022) introduces an alternative “anhydrobiotic model” suggesting that the primary role of fungus-acquired carbon, particularly in the form of polyols, is the ability to survive anabiosis under repeated desiccation. This is particularly important for fungi that exposed the hyphae to aerophytic conditions on the substrate surface. The anhydrobiotic model is based on three key findings: (1) the amount of carbon fixed by a phototroph is 10–20 × greater than is needed to supply the growth and respiratory requirements of the slowly growing fungus (Smith, 1980); (2) mass loss of algal and fungal polyols during repeated re-wetting (Farrar, 1976a); (3) little incorporation of fixed carbon in proteins (Farrar, 1976b). This model was already outlined by (Smith, 1979) who concluded that the symbiotic polyol transfer represents primarily an adaptation to cyclical desiccation and rewetting.

The average ^13^C enrichment of the symbiotic system is lower in semilichens and green biofilms than in lichens (**Fig. 8E**) as is the flux of carbon from algal polyols and sugars to fungal ones (**Fig. 7** vs. **Fig. 8A-D**). Thus, a nutritional model is unlikely for semilichen mycobiont, which tends to reduce its biomass to a minimum, whereas an anhydrobiotic model, where the mycobiont uses a common polyol pool specifically to survive repeated desiccation, seems more likely.

### Aerophytic algae are frequently associated with fungi

According to our observations, aerophytic algae occurring for a long time under conditions of frequent alternation of moisture and desiccation are associated with fungi. Sometimes algae are parasitized by fungi, e.g. by *Athelia* (Motiejūnaitė & Jucevicienė, 2005), but more often it is a longer lasting relationship in which the algae thrive. In general, aerophytic algae occur in some of these symbiotic systems: green biofilms, lichens or semilichens. Algal colonies with no apparent fungal presence were observed only under balanced microclimate conditions with permanently raised humidity. In a periodically drying environment, the algae either need a fungal partner, or, at least, find coexistence with the fungus advantageous (see data in Freystein & Reisser, 2010). During cyclic desiccation and re-hydration, aerophytic algae are subjected to photo-oxidative stress (Candotto Carniel *et al*., 2015), which they cope with by various mechanisms (summarized in Gasulla et al., 2021). Aerophytic algae have been shown to be able to survive short-term stress (on the order of tens of days) caused by desiccation (Candotto Carniel *et al*., 2015), but the longer-term survival of anhydrobiosis or repeated alternations of desiccation and wetting has not been properly investigated.

Attention is currently turning to the protective role of the drought-induced non-photochemical quenching (d-NPQ; Kosugi et al., 2009; Wieners et al., 2018). In this context, Kosugi et al. (2013) demonstrated the transport of fungal arabitol into algae in a lichen. Arabitol acquired by the alga is then essential for the expression of d-NPQ to dissipate excess captured light energy into heat, protecting the photobiont from photoinhibition. Interestingly, no other polyol studied (ribitol, mannitol) was able to enhance d-NPQ. Furthermore, algae can utilise fungal respiratory CO_2_ to increase their photosynthesis and reduce photorespiration (ten Veldhuis *et al*., 2020). In addition, recent research has revealed that algae from the Trebouxiophycae, typically included in symbiotic systems with fungi, share genes for the production of a particular enzyme from the glycoside hydrolases group (Puginier *et al*., 2024). This enzyme may be active in breaking down the lichenan, a key component of the mycobiont’s cell wall. These new findings support the view that aerophytic algae are under selection pressure to symbiosis with their fungal partners.

## Acknowledgement

Linda in Arcadia kindly revised the manuscript. This work was supported by a long-term research development grant RVO [grant number 67985939].

## Supplementary figures S1-S8

**Fig. S1. Corticolous green biofilms collected in an urbanised landscape and observed in visible light**. A, B, on bark of *Carpinus betulus*; C, D, *Picea mariana*; E, F, *Pinus sylvestris*. B, D, F, details of algal-fungal associations. Scales, A, C, E, 0.5 mm; B, D, F, 20 µm.

**Fig. S2. Semilichen ‘*Arthopyrenia*’ *salicis* (Dothydeomycetes, Capnodiales)**. A, B, fruiting bodies (perithecia) on the surface of bark of *Corylus avellana*. A, observed in visible light; B, observed with fluorescence in blue light. C, D, details of algal-fungal association with fluorescence in blue light. Scales, A, B, 0.2 mm; C, D, 50 µm.

**Fig. S3. Semilichens, Ostropomycetidae, Ostropales**. A, B, *Cryptodiscus tabularum*, vertical section of a fruiting body and surrounding substrate. A, observed in visible light; B, observed with fluorescence in blue light. C, D, *Karstenia idaei*; C, vertical section of a fruiting body and surrounding substrate; D, detail of the excipulum of fungal fruiting body with algae of the *Coccomyxa*/*Elliptochloris* type. Scales, A-C, 100 µm; D, 20 µm.

**Fig. S4. Semilichen *Cyrtidula quercus* (Dothydeomycetes, incertae sedis) associated with *Trentepohlia* sp**. A, observed in visible light; B, observed with fluorescence in blue light. Scales, 100 µm.

**Fig. S5. Semilichen *Naevia punctiformis* (Arthoniomycetes, Arthoniales)**. A, observed in visible light; B, observed with fluorescence in blue light. Note the many scattered algal cells observable by red chlorophyll autofluorescence. Scales, 0.2 mm.

**Fig. S6. Chlorophyll *a* fluorescence kinetics of green biofilm, semilichen *Arthonia salicis* and true lichen *Graphis scripta* (denoted in pictures)**. Visual situation (upper subplots), absolute values fluorescence (middle subplot) and fluorescence standardised to F_o_ = 1 (bottom suplots). Comparison of symbioses (blue line), plant tissue where photobionts were removed (red line) and calibration plate where no photochemistry occurs (black line) is made in each system. Dark adapted samples were subjected to low intensity measuring light (< 1 µmol m^-2^ s^-1^) for first 4 s, then to saturating flash (≈ 1000 µmol m^-2^ s^-1^) for one second allowing us to calculate maximum quantum yield of PS II (F_v_/F_m_). After short dark relaxation (6 to 19s), in time 20 to 90 s, actinic light (150 µmol m^-2^ s^-1^) with five superimposed saturating flashes were applied to find photochemical (photosynthesis) and non-photochemical (photoprotection) quenching. Last part (91-190s) is dark relaxation with three saturating flashes to obtain relaxation rate of photoprotective mechanisms.

**Fig. S7. Percentage of carbon in particular metabolites of total metabolite pool (left) and percentage of new carbon (**^**13**^**C) in these metabolites 2 h, one day and four days after** ^**13**^**CO**_**2**_ **labelling (right)**. First two pages represent corticolous green biofilm and seven semilichens, last page are four control true lichens. Gradual incorporation of new carbon into fungal polyols (arabitol and mannitol) is visible.

**Fig S8. Average** ^**13**^**C enrichment in all metabolites pooled 2 h, one day and four days after** ^**13**^**CO**_**2**_ **labelling**. It is measure of CO_2_ assimilation intensity (and reciprocally metabolite turnover) for whole symbiotic system. We can see, true lichens (blue lines) are typically the most enriched despite large proportion of heterotrophic (fungal) biomass. Corticolous green biofilm (green line) is intermediate but only one system measured is not sufficient to estimate mean and variability in these systems. Finally, semilichens (red lines) are less but substantially enriched. Note: y-axis is semi logarithmic.

## Notes

### Competing Interest Statement

The authors have declared no competing interest.

### Summary of Updates

Symbiotic systems of photosynthetic microorganisms and fungi are widespread in terrestrial biomes and lichens are probably the most advanced and complex. Conversely, the least complex systems are "green biofilms" with a completely unexplored mycobiome. We describe here a new system intermediate between green biofilms and lichens: semilichens. Light and fluorescence microscopy, eDNA sequencing, molecular phylogeny, Chlorophyll a fluorescence and 13C labelling/metabolomics were used to study algal and fungal identity, morphology and physiology of the symbiosis. Tight contact between algae and a single predominant fungus (mycobiont) is revealed in semilichens. The algae are from the symbiotic lineages of Trebouxiophyceae and Ulvophyceae, the fungi belong to Arthoniomycetes, Dothideomycetes, Eurotiomycetes, Lecanoromycetes and Lichinomycetes. Algae are alive and perform substantial photosynthetic activity. 13C labelled photosynthates are partially converted into specific fungal polyols (arabitol, mannitol) demonstrating the C-flow from algae to fungi. The new symbiotic system was defined and compared with other terrestrial algal-fungal symbioses. It is characterized by minimalist environmental requirements and extremely low production of biomass. As a result, it also inhabits environments unfavourable for lichens. Our research supports the hypothesis that the long-term existence of algae and fungi in terrestrial conditions affected by frequent and repeated drying is likely dependent on their mutual coexistence.

## References

Andrews S. 2010. FastQC: a quality control tool for high throughput sequence data. https://www.bioinformatics.babraham.ac.uk/projects/fastqc/.

Aras S, Cansaran D. 2006. Isolation of DNA for sequence analysis from herbarium material of some lichen specimens. Turkish Journal of Botany 30: 449–453.

Bálint M, Schmidt PA, Sharma R, Thines M, Schmitt I. 2014. An Illumina metabarcoding pipeline for fungi. Ecology and Evolution 4:2642–2653.

Beckett R, Ntombela N, Scott E, Gurjanov O, Minibayeva F, Liers C. 2015. Role of laccases and peroxidases in saprotrophic activities in the lichen Usnea undulata. Fungal Ecology 14:71–78.

Bonito G. 2024. Ecology and evolution of algal–fungal symbioses. Current Opinion in Microbiology 79.

Candotto Carniel F, Zanelli D, Bertuzzi S, Tretiach M. 2015. Desiccation tolerance and lichenization: a case study with the aeroterrestrial microalga Trebouxia sp. (Chlorophyta). Planta 242:493–505.

Chen C-C, Xie Q-Y, Chuang P-S, Darnajoux R, Chien Y-Y, Wang W-H, Tian X, Tu C-H, Chen B-C, Tang S-L, et al. 2025. A thallus-forming N-fixing fungus-cyanobacterium symbiosis from subtropical forests.

Díaz-Escandón D, Tagirdzhanova G, Vanderpool D, Allen CCG, Aptroot A, Češka O, Hawksworth DL, Huereca A, Knudsen K, Kocourková J, et al. 2022. Genome-level analyses resolve an ancient lineage of symbiotic ascomycetes. Current Biology 32:5209-5218.e5.

Drew E, Smith D. 1967. Studies in the physiology of lichens. VIII.Movement of glucose from alga to fungus during photosynthesis in thallus of Peltigera polydactyla. New Phytologist 66:389–400.

Ettl H, Gärtner G. 1995. Syllabus der Boden-, Luft- und Flechtenalgen. Stuttgart/Jena/New York: Gustav Fischer Verlag.

Farrar J. 1976a. Ecological physiology of the lichen Hypogymnia physodes. II. Effects of wetting and drying cycles and the concept of ‘physiological buffering’. New Phytologist 77:105–113.

Farrar J. 1976b. Ecological physiology of the lichen Hypogymnia physodes. I. Some effects of constant water saturation. New Phytologist 77:93–103.

Feige G, Kremer B. 1980. Unusual carbohydrate pattern in Trentepohlia species. Phytochemistry: 1844–1845.

Freudenthal J, Dumack K, Schaffer S, Schlegel M, Bonkowski M. 2024. Algal-fungi symbioses and bacteria-fungi co-exclusion drive tree species-specific differences in canopy bark microbiomes. ISME Journal, wrae206.

Freystein K, Reisser W. 2010. Green Biofilms on Tree Barks: More than Just Algae. In: Seckbach J, Grube M, eds. Symbioses and Stress. Springer, 557–573.

Gärtner G. 1974. Beitrag zur Systematik und Ökologie von Rindenalgen.

Gasulla F, Del Campo EM, Casano LM, Guéra A. 2021. Advances in understanding of desiccation tolerance of lichens and lichen-forming algae. Plants 10.

Geitler L. 1942. Morphologie, Entwicklungsgeschichte und Systematik neuerer bemerkenswerter aerophytischer Algen aus Wien. Flora 136:1–29.

Gilbert O. 1971. Studies along edge of a lichen desert. Lichenologist 5:11–17.

Gustavs L, Eggert A, Michalik D, Karsten U. 2010. Physiological and biochemical responses of green microalgae from different habitats to osmotic and matric stress. Protoplasma 243:3–14.

Gustavs L, Görs M, Karsten U. 2011. Polyol patterns in biofilm-forming aeroterrestrial green algae (Trebouxiophyceae, Chlorophyta). Journal of Phycology 47:533–537.

Honegger R. 1993. Tansley Review No. 60 Developmental biology of lichens. New Phytologist 125:659–677.

Kohlmeyer J, Hawksworth DH, Volkmann-Kohlmeyer B. 2004. Observations on two marine and maritime “borderline” lichens: Mastodia tessellata and Collemopsidium pelvetiae. Mycological Progress 3:51–56.

Kosugi M, Arita M, Shizuma R, Moriyama Y, Kashino Y, Koike H, Satoh K. 2009. Responses to desiccation stress in lichens are different from those in their photobionts. Plant and Cell Physiology 50:879–888.

Kosugi M, Miyake H, Yamakawa H, Shibata Y, Miyazawa A, Sugimura T, Satoh K, Itoh S, Kashino Y. 2013. Arabitol provided by lichenous fungi enhances ability to dissipate excess light energy in a symbiotic green alga under desiccation. Plant and Cell Physiology 54:1316–1325.

Kubásek J, Hájek T, Duckett J, Pressel S, Šantrůček J. 2021. Moss stomata do not respond to light and CO2 concentration but facilitate carbon uptake by sporophytes: a gas exchange, stomatal aperture, and 13C-labelling study. New Phytologist 230:1815–1828.

Kulichová J, Škaloud P, Neustupa J. 2014. Molecular diversity of green corticolous microalgae from two sub-Mediterranean European localities. European Journal of Phycology 49:345–355.

Laufer Z, Beckett RP, Minibayeva F V, Lüthje S, Böttger M. 2009. Diversity of laccases from lichens in suborder Peltigerineae.

Lewis DH, Smith DC. 1967. Sugar alcohols (polyols) in fungi and green plants I. Distribution, physiology and metabolism. New Phytologist 66:143–184.

Lücking R, Leavitt SD, Hawksworth DL. 2021. Species in lichen-forming fungi: balancing between conceptual and practical considerations, and between phenotype and phylogenomics. Fungal Diversity 109:99–154.

Lücking R, Lumbsch HT, D. Stéfano JF, Lizano D, Carranza J, Bernecker A, Chaves JL, Umaña L, Facio R, Pedro S, et al. 2008. Eremithallus costaricensis (Ascomycota: Lichinomycetes: Eremithallales), a new fungal lineage with a novel lichen symbiotic lifestyle discovered in an urban relict forest in Costa Rica. Symbiosis 46:161–170.

Lücking R, Spribille T. 2024. The Lives of Lichens: A Natural History. Princeton, Oxford: Princeton University Press.

Mahé F, Rognes T, Quince C, de Vargas C, Dunthorn M. 2014. Swarm: Robust and fast clustering method for amplicon-based studies. PeerJ 2014.

Motiejūnaitė J, Jucevicienė N. 2005. Epidemiology of the fungus Athelia arachnoidea in epiphytic communities of broadleaved forests under strong anthropogenic impact. Ekologija 4:28–34.

Muggia L, Baloch E, Stabentheiner E, Grube M, Wedin M. 2011. Photobiont association and genetic diversity of the optionally lichenized fungus Schizoxylon albescens. FEMS Microbiology Ecology 75:255–272.

Nguyen LT, Schmidt HA, Von Haeseler A, Minh BQ. 2015. IQ-TREE: A fast and effective stochastic algorithm for estimating maximum-likelihood phylogenies. Molecular Biology and Evolution 32:268– 274.

Palmqvist K, Dahlman L, Jonsson A, Nash TH III. 2008. The carbon economy of lichens. In: Nash TI, ed. Lichen biology, 2nd edition. Cambridge, UK: Cambridge University Press, 184–217.

Pérez-Ortega S, Garrido-Benavent I, Grube M, Olmo R, de los Ríos A. 2016. Hidden diversity of marine borderline lichens and a new order of fungi: Collemopsidiales (Dothideomyceta). Fungal Diversity 80:285–300.

Pichler G, Muggia L, Carniel FC, Grube M, Kranner I. 2023. How to build a lichen: from metabolite release to symbiotic interplay. New Phytologist 238:1362–1378.

Puginier C, Libourel C, Otte J, Skaloud P, Haon M, Grisel S, Petersen M, Berrin JG, Delaux PM, Dal Grande F, et al. 2024. Phylogenomics reveals the evolutionary origins of lichenization in chlorophyte algae. Nature Communications 15.

Richardson DHS, Smith DC. 1968. Lichen physiology: IX. Carbohydrate movement from The Trebouxia symbiobr of Xanthoria aureola to the fungus. New Phytologist 67:61–68.

Richardson D, Smith D, Lewis D. 1967. Carbohydrate movement between the symbionts of lichens. Nature 214:879–882.

Schwendener S. 1869. Die Algentypen der Flechtengonidien. Basel, Switzerland: Universitaetsbuchdruckerei.

Da Silva Cáceres ME, Lücking R, Schumm F, Aptroot A. 2020. A lichenized family yields another renegade lineage: Papilionovela albothallina is the first non-lichenized, saprobic member of Graphidaceae subfam. Graphidoideae. Bryologist 123:144–154.

Smith D. 1963. Studies in the physiology of lichens. IV. Carbohydrates in Peltigera polydactyla and the utilization of absorbed glucose. New Phytologist 62:205–216.

Smith D. 1979. Is a lichen a good model of interactions in nutrient-limited environments? In: Shilo M, ed. Strategies of microbial life in extreme environments: report of the Dahlem Workshop on Strategy of Life in Extreme Environments, Berlin, 1978, November 20–24. Weinheim, Germany: Verlag Chemie, 291–301.

Smith D. 1980. Mechanisms of nutrient movement between the lichen symbionts. In: Cook C, Pappas P, Rudolph E, eds. Cellular interactions in symbiosis and parasitism. Columbus, OH, USA: Ohio State University Press, 197–221.

Spribille T, Resl P, Stanton DE, Tagirdzhanova G. 2022. Evolutionary biology of lichen symbioses. New Phytologist 234:1566–1582.

Štifterová A, Neustupa J. 2015. Community structure of corticolous microalgae within a single forest stand: Evaluating the effects of bark surface ph and tree species. Fottea 15:113–122.

Štifterová A, Neustupa J. 2017. Small-scale variation of corticolous microalgal covers: Effects of microhabitat, season, and space. Phycological Research 65:299–311.

Thiyagaraja V, Lücking R, Ertz D, Wanasinghe DN, Karunarathna SC, Camporesi E, Hyde KD. 2020. Evolution of non-lichenized, saprotrophic species of Arthonia (Ascomycota, Arthoniales) and resurrection of Naevia, with notes on Mycoporum. Fungal Diversity 102:205–224.

ten Veldhuis MC, Ananyev G, Dismukes GC. 2020. Symbiosis extended: exchange of photosynthetic O2 and fungal-respired CO2 mutually power metabolism of lichen symbionts. Photosynthesis Research 143:287–299.

Větrovský T, Baldrian P, Morais D. 2018. SEED 2:A user-friendly platform for amplicon high-throughput sequencing data analyses. In: Bioinformatics. Oxford University Press, 2292–2294.

Vondrák J, Svoboda S, Malíček J, Palice Z, Kocourková J, Knudsen K, Mayrhofer H, Thüs H, Schultz M, Košnar J, et al. 2022. From Cinderella to Princess: An exceptional hotspot of lichen diversity in a long-inhabited central-European landscape. Preslia 94:143–181.

Vondrák J, Svoboda S, Zíbarová L, Štenclová L, Mareš J, Pouska V, Košnar J, Kubásek J. 2023. Alcobiosis, an algal-fungal association on the threshold of lichenisation. Scientific Reports 13.

Wedin M, Döring H, Gilenstam G. 2004. Saprotrophy and lichenization as options for the same fungal species on different substrata: Environmental plasticity and fungal lifestyles in the Stictis-Conotrema complex. New Phytologist 164:459–465.

Wieners PC, Mudimu O, Bilger W. 2018. Survey of the occurrence of desiccation-induced quenching of basal fluorescence in 28 species of green microalgae. Planta 248:601–612.

Wijayawardene NN, Hyde KD, Al-Ani LKT, Tedersoo L, Haelewaters D, Rajeshkumar KC, Zhao RL, Aptroot A, Leontyev D V., Saxena RK, et al. 2020. Outline of Fungi and fungus-like taxa. Mycosphere 11:1060–1456.

Zukal H. 1891. Halbflechten. Flora (Marburg) 74:92–107.

