## Supplementary file for "Semilichen an unjustly neglected symbiotic system between green biofilms and true lichens"

**Fig. S1. Corticolous green biofilms collected in an urbanised landscape and observed in visible light.** A, B, on bark of *Carpinus betulus*; C, D, *Picea mariana*; E, F, *Pinus sylvestris*. B, D, F, details of algal-fungal associations. Scales, A, C, E, 0.5 mm; B, D, F, 20 µm.

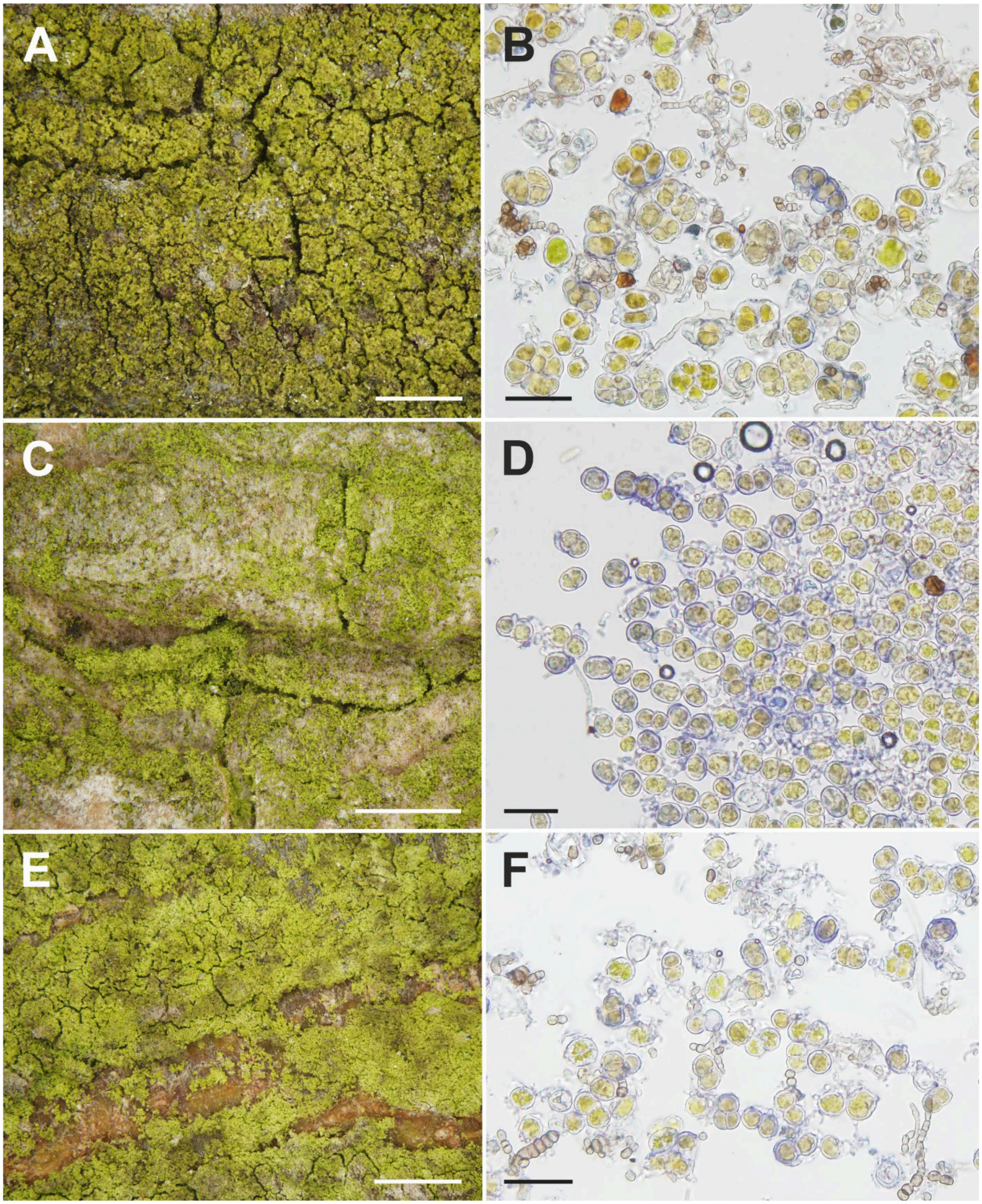

**Fig. S2. Semilichen '*Arthopyrenia*' *salicis* (Dothydeomycetes, Capnodiales).** A, B, fruiting bodies (perithecia) on the surface of bark of *Corylus avellana*. A, observed in visible light; B, observed with fluorescence in blue light. C, D, details of algal-fungal association with fluorescence in blue light. Scales, A, B, 0.2 mm; C, D, 50 µm.

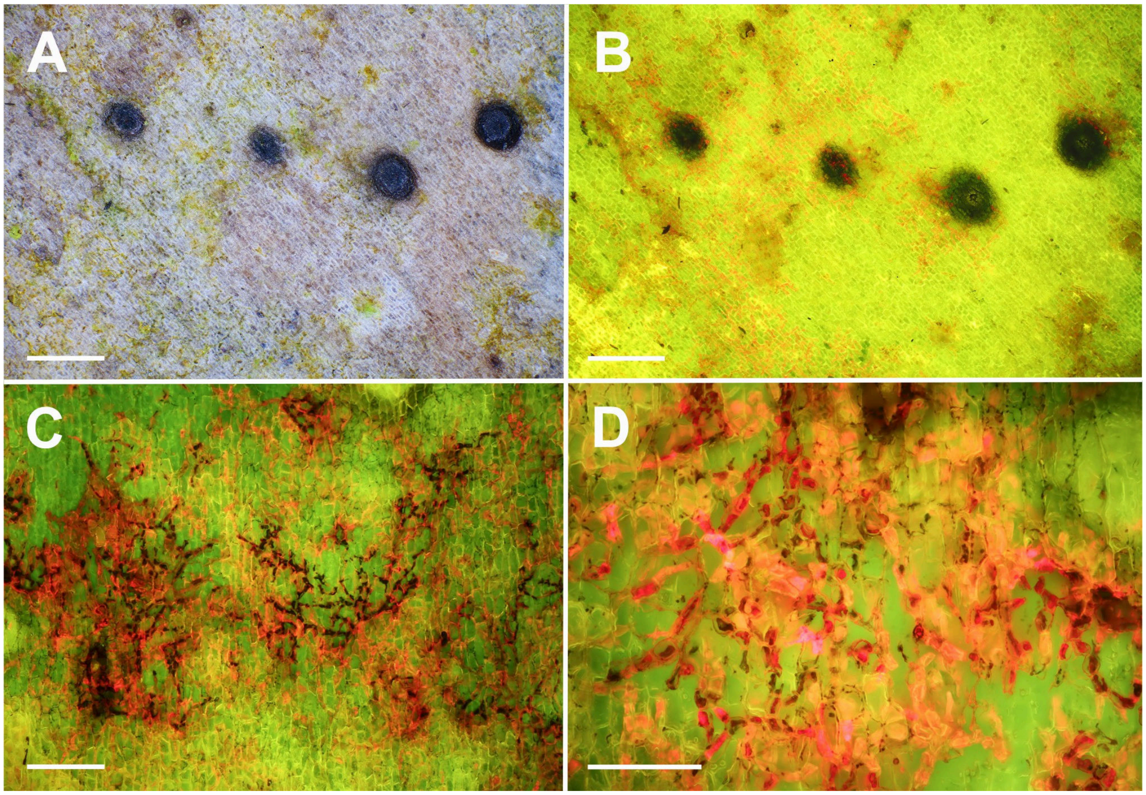

**Fig. S3. Semilichens, Ostropomycetidae, Ostropales.** A, B, *Cryptodiscus tabularum*, vertical section of a fruiting body and surrounding substrate. A, observed in visible light; B, observed with fluorescence in blue light. C, D, *Karstenia idaei*; C, vertical section of a fruiting body and surrounding substrate; D, detail of the excipulum of fungal fruiting body with algae of the *Coccomyxa*/*Elliptochloris* type. Scales, A-C, 100 µm; D, 20 µm.

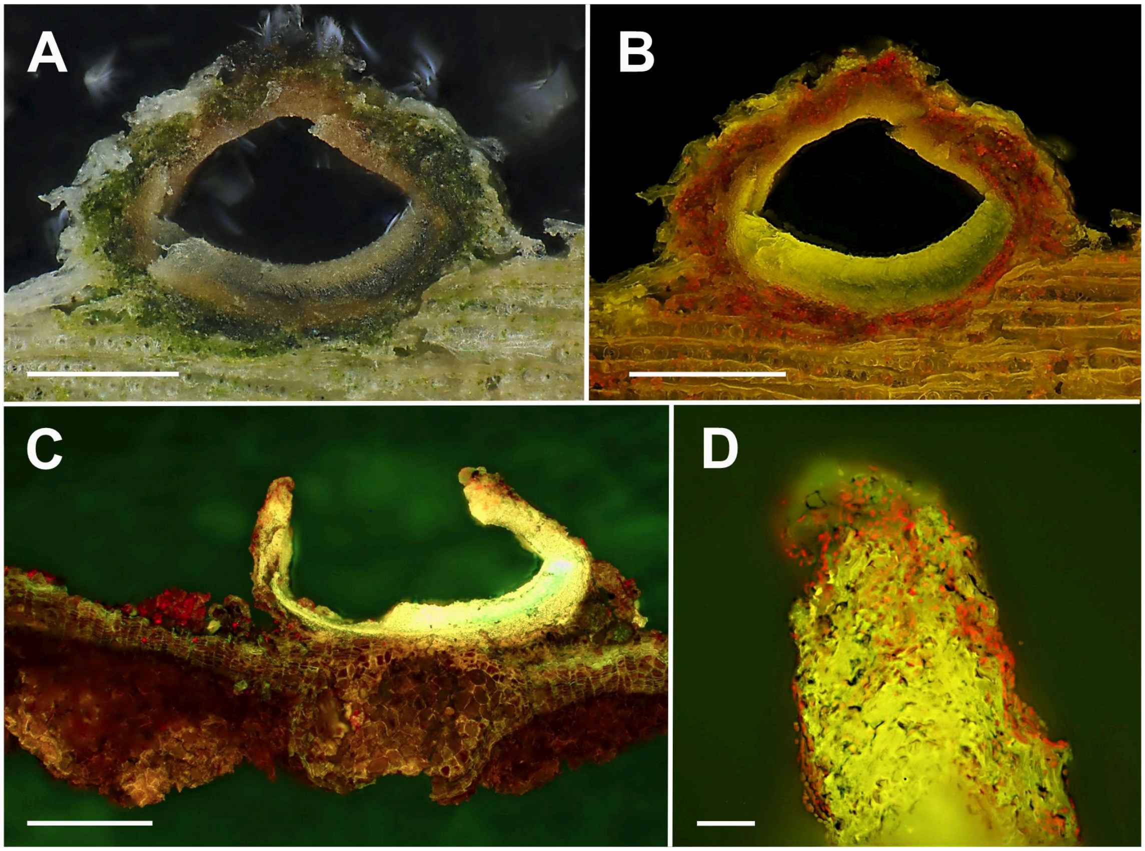

**Fig. S4. Semilichen *Cyrtidula quercus* (Dothydeomycetes, incertae sedis) associated with *Trentepohlia* sp.** A, observed in visible light; B, observed with fluorescence in blue light. Scales, 100 µm.

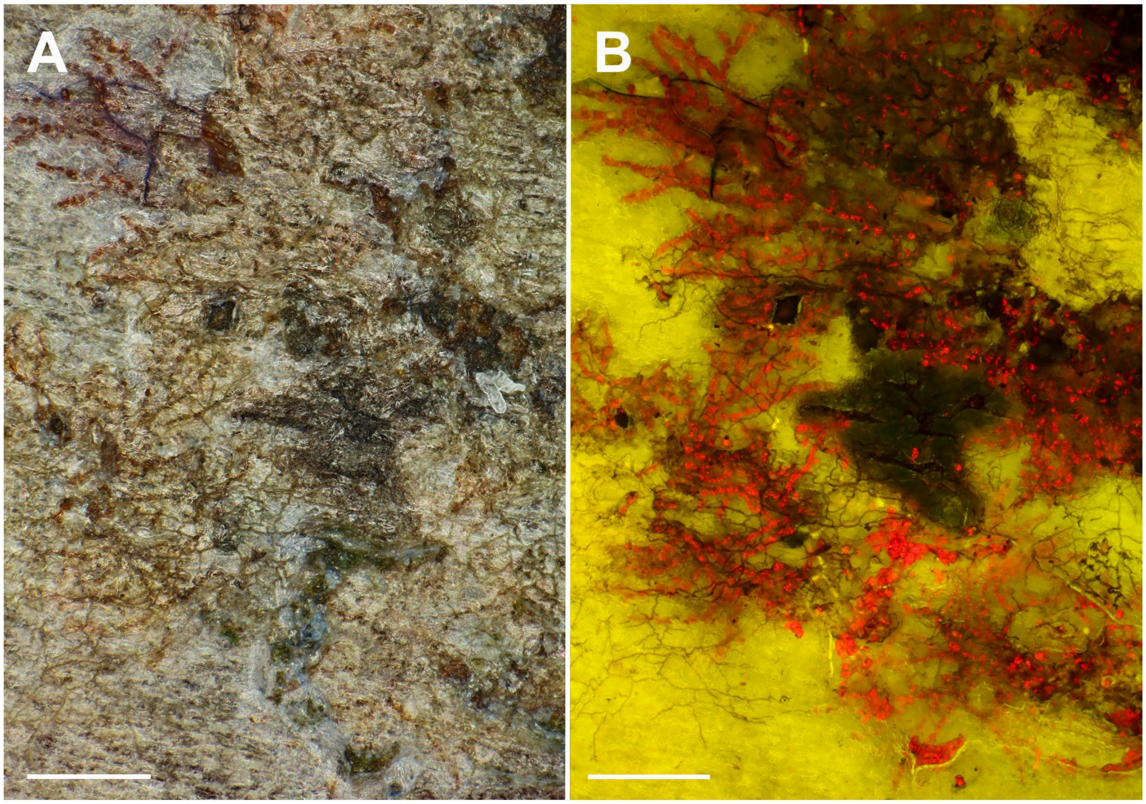

**Fig. S5. Semilichen *Naevia punctiformis* (Arthoniomycetes, Arthoniales).** A, observed in visible light; B, observed with fluorescence in blue light. Note the many scattered algal cells observable by red chlorophyll autofluorescence. Scales, 0.2 mm.

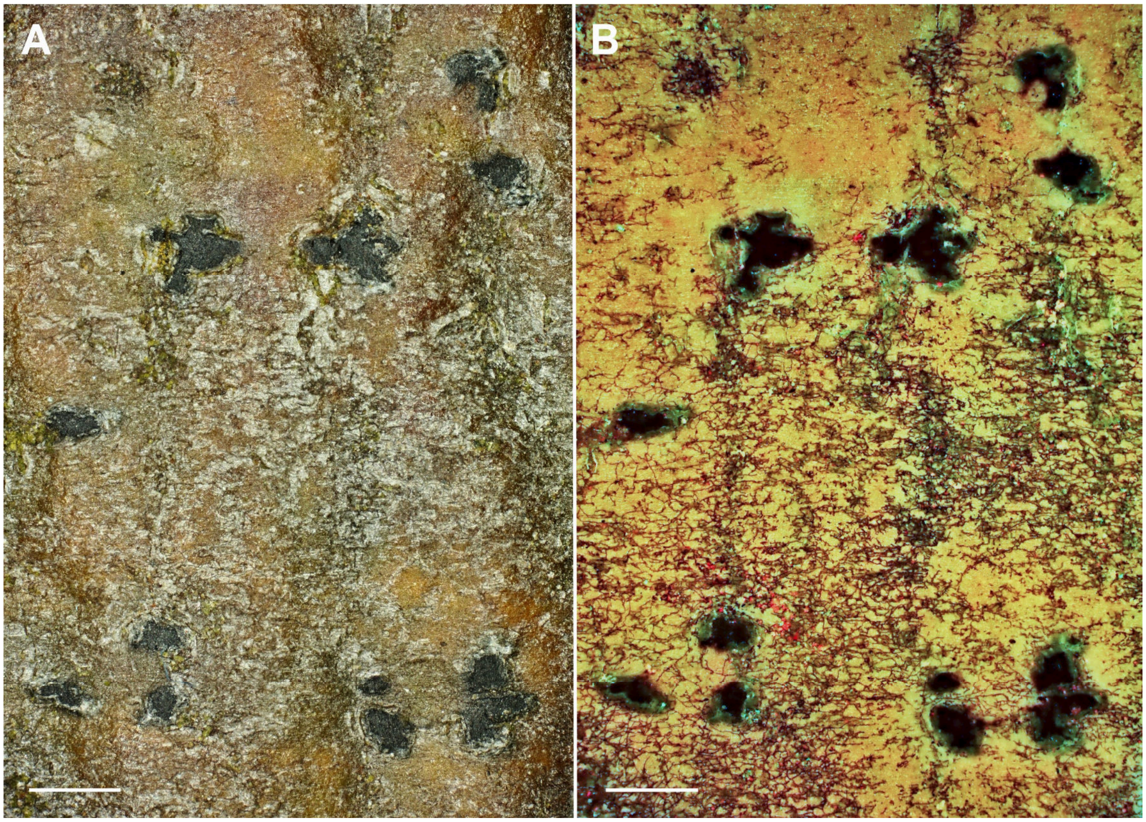

**Fig. S6. Chlorophyll *a* fluorescence kinetics of green biofilm, semilichen *Arthonia salicis* and true lichen *Graphis scripta* (denoted in pictures).** Visual situation (upper subplots), absolute values fluorescence (middle subplot) and fluorescence standardised to $F_o = 1$ (bottom suplots). Comparison of symbioses (blue line), plant tissue where photobionts were removed (red line) and calibration plate where no photochemistry occurs (black line) is made in each system. Dark adapted samples were subjected to low intensity measuring light (< 1 µmol $m^{-2}$ $s^{-1}$) for first 4 s, then to saturating flash (≈ 1000 µmol $m^{-2}$ $s^{-1}$ ) for one second allowing us to calculate maximum quantum yield of PS II ($F_v/F_m$). After short dark relaxation (6 to 19s), in time 20 to 90 s, actinic light (150 µmol $m^{-2}$ $s^{-1}$) with five superimposed saturating flashes were applied to find photochemical (photosynthesis) and non-photochemical (photoprotection) quenching. Last part (91-190s) is dark relaxation with three saturating flashes to obtain relaxation rate of photoprotective mechanisms.

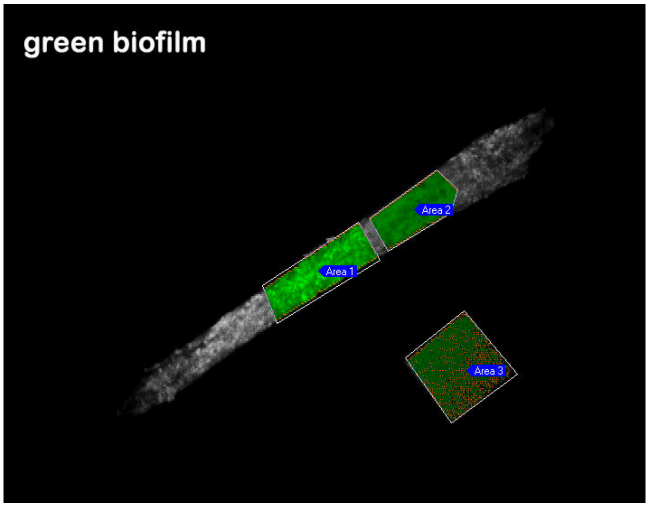
green biofilm
Area 1
Area 2
Area 3

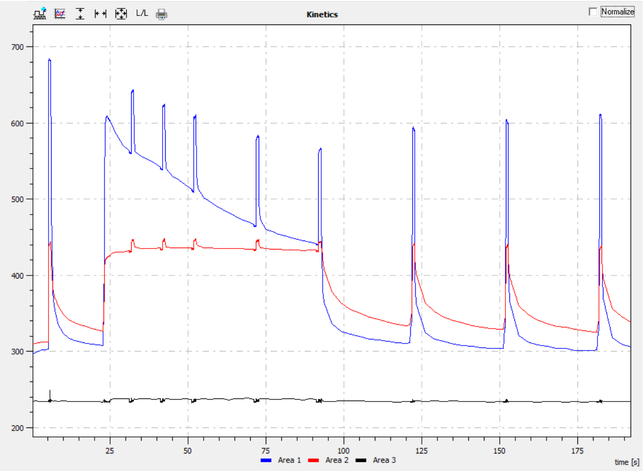
Kinetics
Normalize
700
600
500
400
300
200
25
50
75
100
125
150
175
time [s]
Area 1
Area 2
Area 3

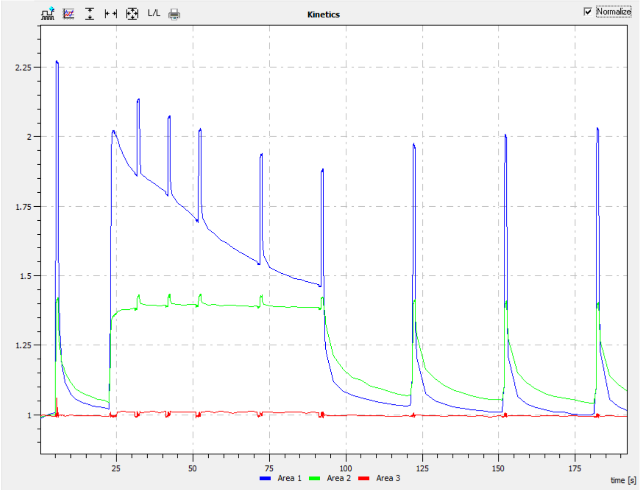
Kinetics
Normalize
2.25
2
1.75
1.5
1.25
1
25
50
75
100
125
150
175
time [s]
Area 1
Area 2
Area 3

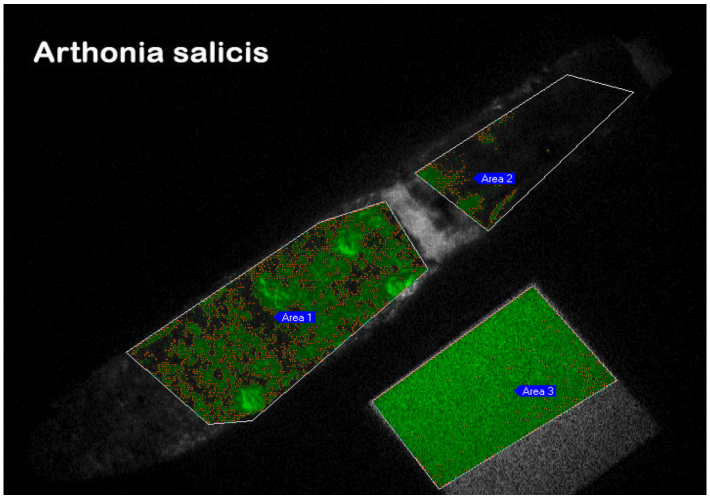
Arthonia salicis
Area 1
Area 2
Area 3

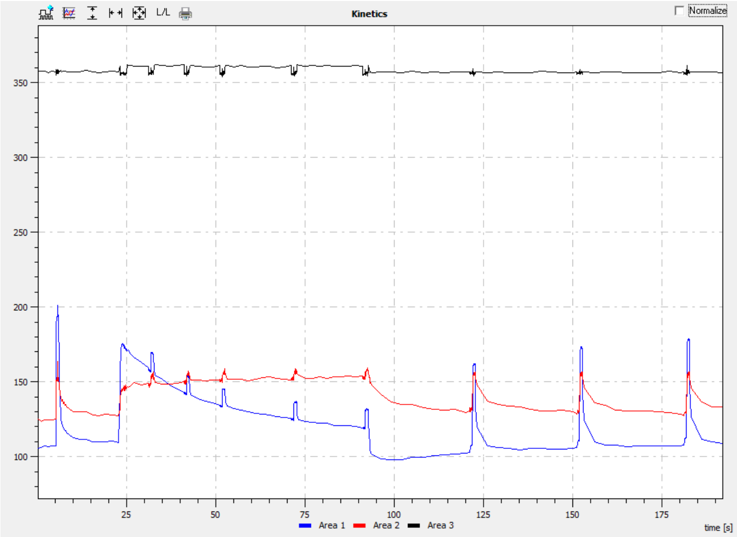
Kinetics
Normalize
350
300
250
200
150
100
25
50
75
100
125
150
175
Area 1
Area 2
Area 3
time [s]

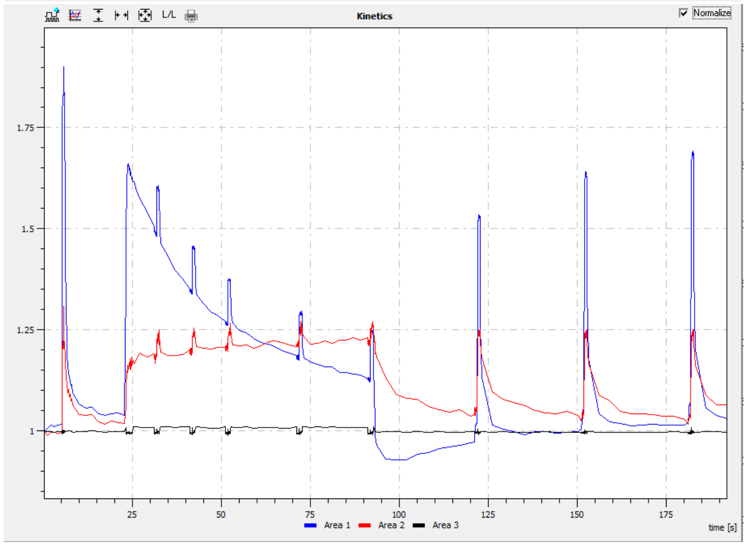
Kinetics
Normalize
1.75
1.5
1.25
1
25
50
75
100
125
150
175
Area 1
Area 2
Area 3
time [s]

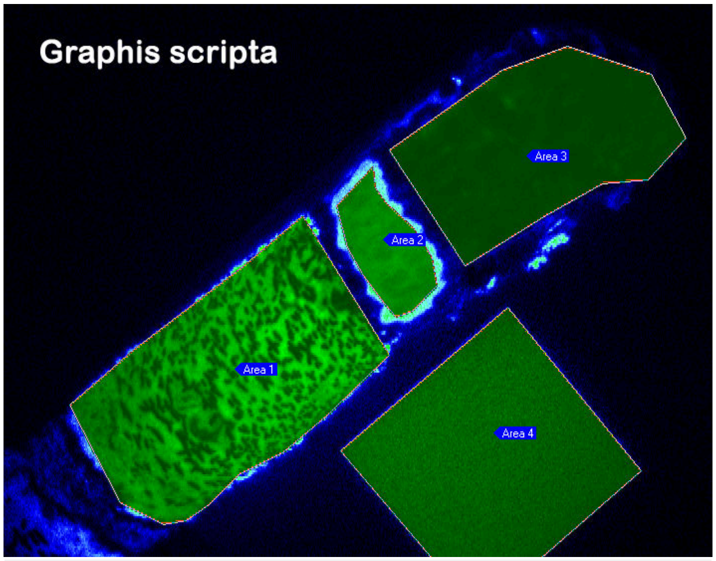
Graphis scripta
Area 1
Area 2
Area 3
Area 4

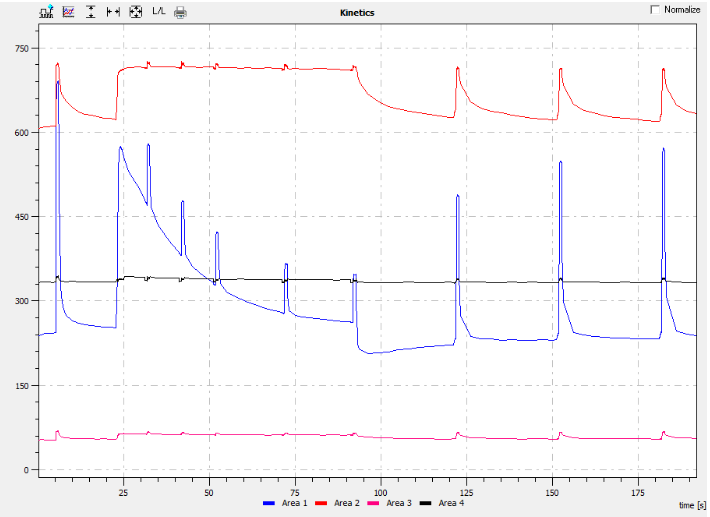
L/L
Kinetics
Normalize
750
600
450
300
150
0
25
50
75
100
125
150
175
time [s]
Area 1
Area 2
Area 3
Area 4

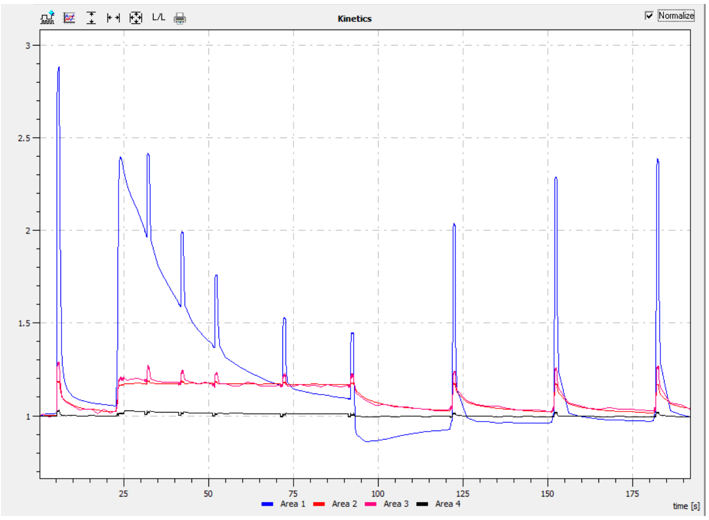
L/L
Kinetics
Normalize
3
2.5
2
1.5
1
25
50
75
100
125
150
175
time [s]
Area 1
Area 2
Area 3
Area 4

**Fig. S7. Percentage of carbon in particular metabolites of total metabolite pool (left) and percentage of new carbon ($^{13}C$) in these metabolites 2 h, one day and four days after $^{13}CO_2$ labelling (right).** First two pages represent corticolous green biofilm and seven semilichens, last page are four control true lichens. Gradual incorporation of new carbon into fungal polyols (arabitol and mannitol) is visible.

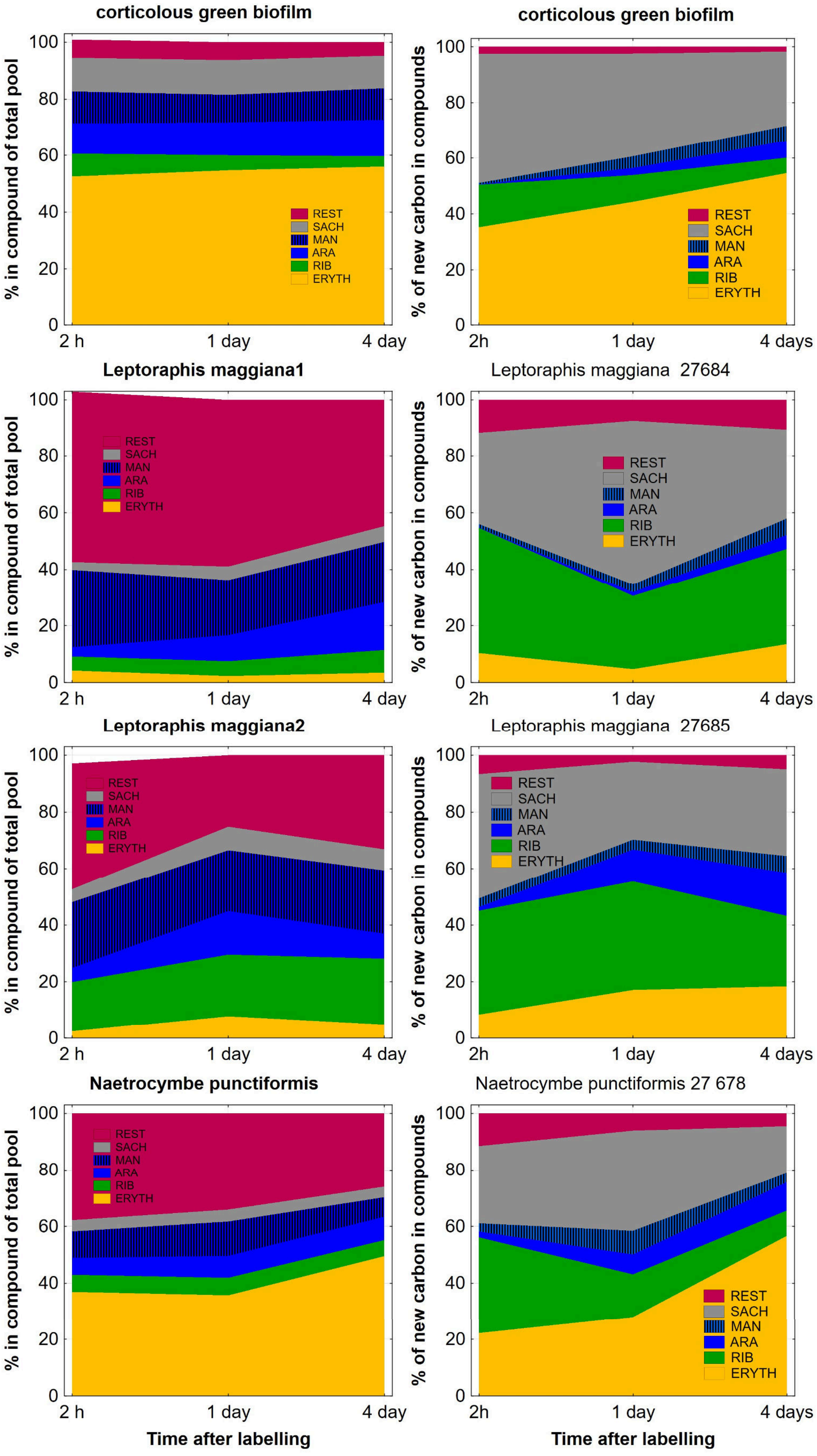
corticolous green biofilm
corticolous green biofilm
Leptoraphis maggiana1
Leptoraphis maggiana 27684
Leptoraphis maggiana2
Leptoraphis maggiana 27685
Naetrocymbe punctiformis
Naetrocymbe punctiformis 27 678
% in compound of total pool
% of new carbon in compounds
REST
SACH
MAN
ARA
RIB
ERYTH
0
20
40
60
80
100
2 h
1 day
4 day
2h
4 days
Time after labelling

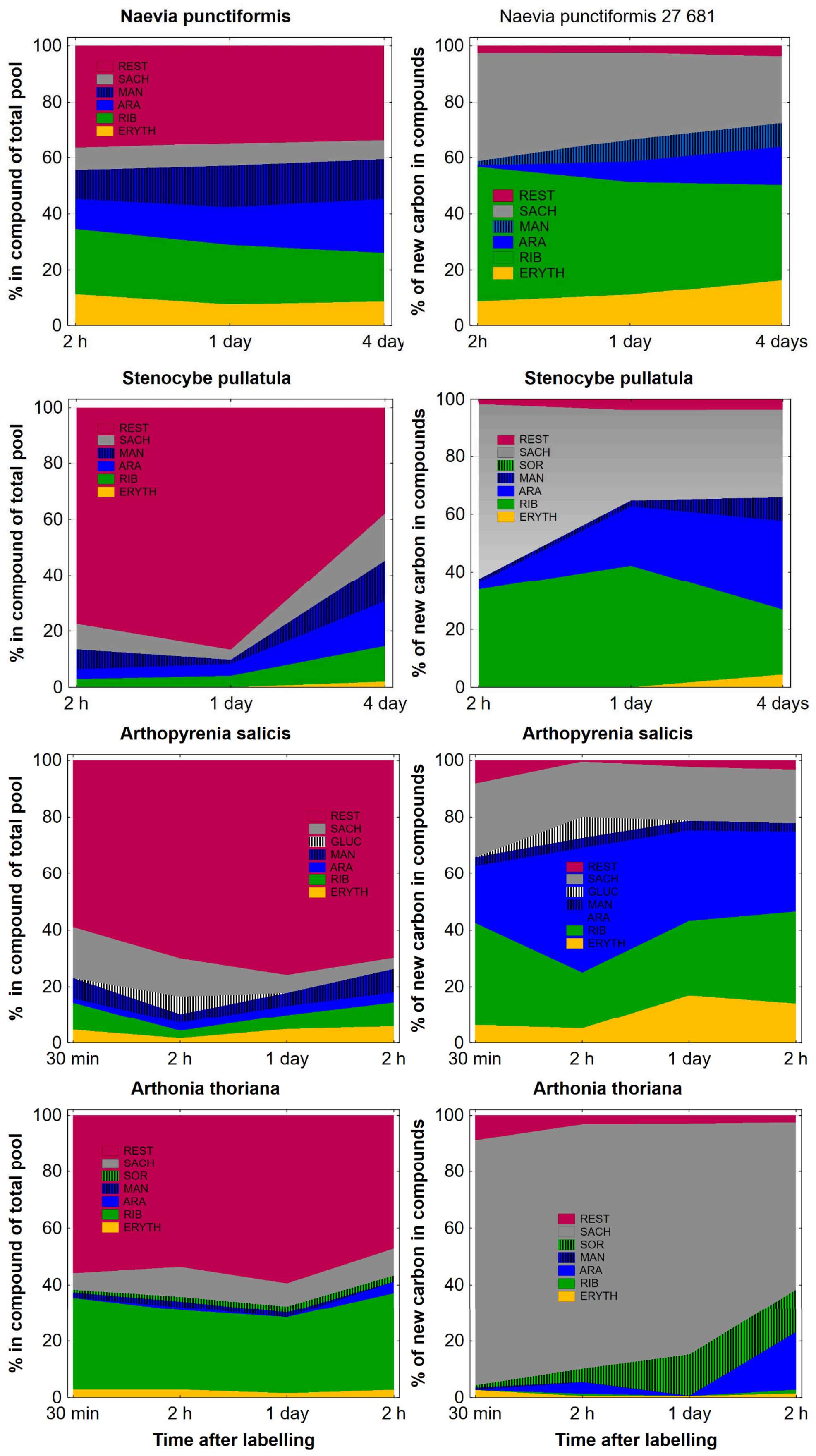

Naevia punctiformis
Naevia punctiformis 27 681
Stenocybe pullatula
Stenocybe pullatula
Arthopyrenia salicis
Arthopyrenia salicis
Arthonia thoriana
Arthonia thoriana
% in compound of total pool
% of new carbon in compounds
REST
SACH
SOR
GLUC
MAN
ARA
RIB
ERYTH
0
20
40
60
80
100
2 h
1 day
4 day
2h
4 days
30 min
Time after labelling

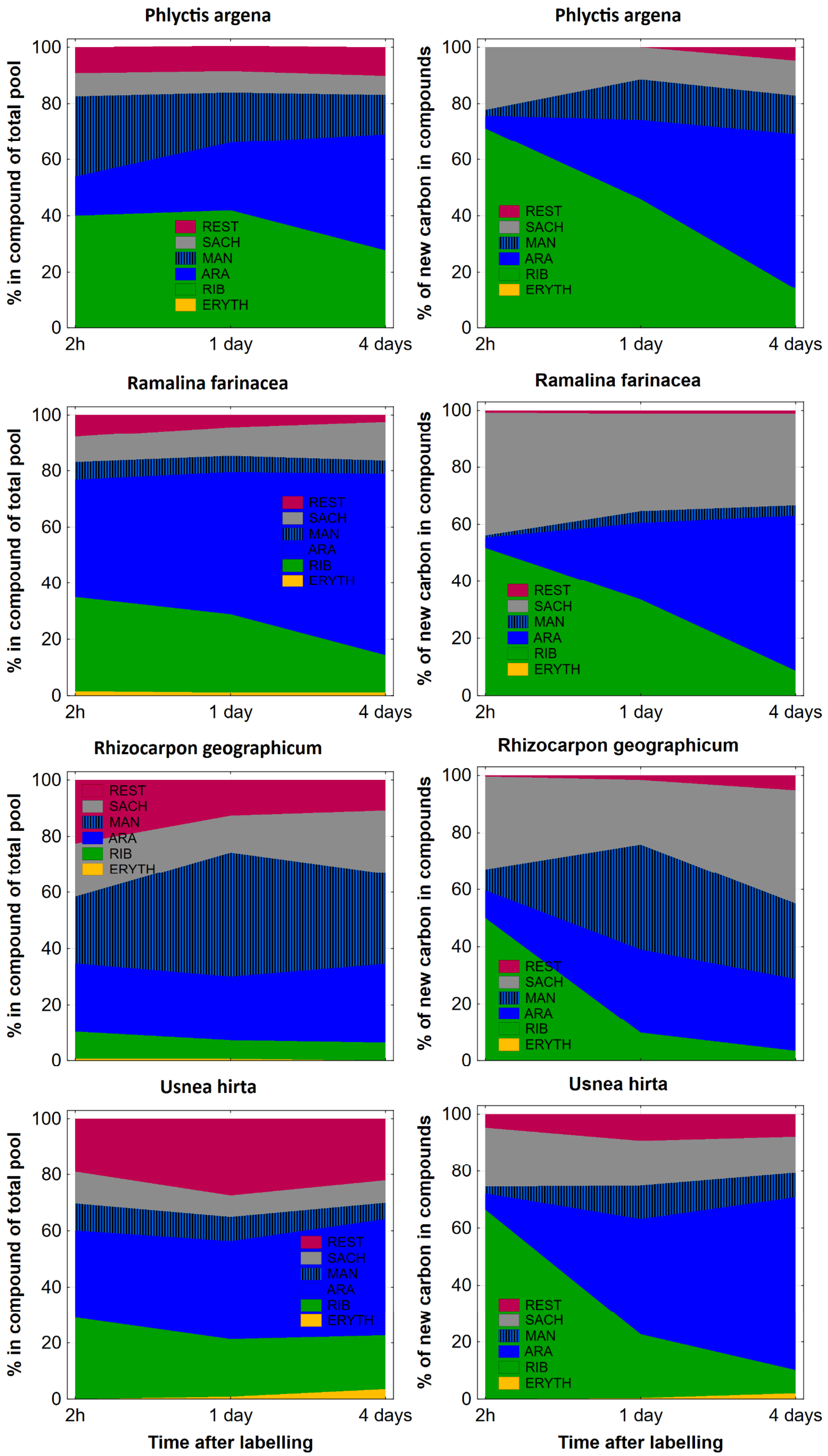

Phlyctis argena
Phlyctis argena
Ramalina farinacea
Ramalina farinacea
Rhizocarpon geographicum
Rhizocarpon geographicum
Usnea hirta
Usnea hirta
% in compound of total pool
% of new carbon in compounds
REST
SACH
MAN
ARA
RIB
ERYTH
0
20
40
60
80
100
2h
1 day
4 days
Time after labelling
Time after labelling

**Fig S8. Average $^{13}C$ enrichment in all metabolites pooled 2 h, one day and four days after $^{13}CO_2$ labelling.** It is measure of $CO_2$ assimilation intensity (and reciprocally metabolite turnover) for whole symbiotic system. We can see, true lichens (blue lines) are typically the most enriched despite large proportion of heterotrophic (fungal) biomass. Corticolous green biofilm (green line) is intermediate but only one system measured is not sufficient to estimate mean and variability in these systems. Finally, semilichens (red lines) are less but significantly enriched. Note: y-axis is semi-logarithmic.

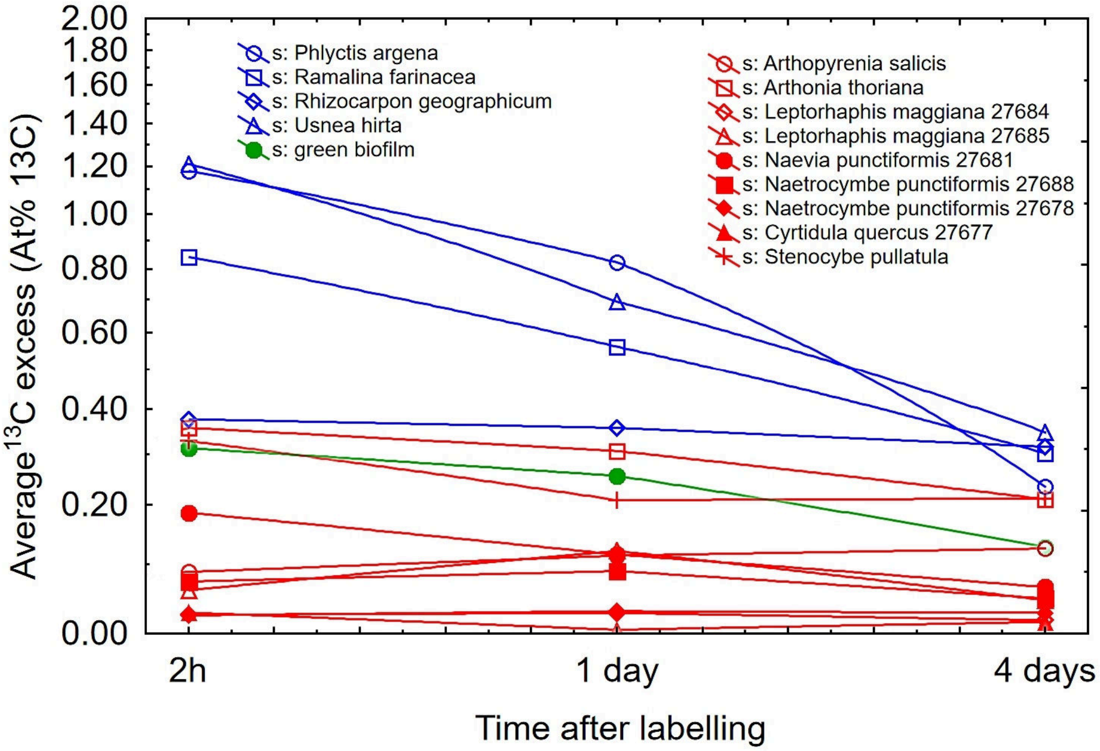
